# Mechanical regulation of cuboidal-to-squamous epithelial transition in the *Drosophila* developing wing

**DOI:** 10.1101/2024.09.21.614249

**Authors:** Stefan Harmansa, Thomas Lecuit

## Abstract

Growing tissues are constantly exposed to mechanical stresses that lead to deformations at the cell and tissue scale. In epithelial organs, cells form monolayers whose thickness can change dramatically during development. Here, we address how cell shape changes in the peripodial epithelium of the *Drosophila* wing disc emerge from the interplay of basement membrane properties with tissue-extrinsic mechanical stress. We show that tissue-extrinsic stress arising from disc proper bending elastically deforms central peripodial cells and induces a cuboidal-to-squamous epithelial transition. In contrast, a rigid basement membrane shields peripheral hinge cells from this bending stress and causes a cuboidal-to-columnar transition. These inverse shape transitions are further amplified by selective shearing of central cells due to coupling via the apical extracellular matrix protein Dumpy. These findings point to a pivotal role of the basement membrane and inter-tissue coupling in the emergence of stress patterns and cell deformations during organ growth.

## Introduction

The process of form generation, known as morphogenesis, entails a complex sequence of 3D deformations that mold growing tissues into functional shapes in embryos and organs^1^. Deformations arise from forces at the cellular and tissue scale. While cells are active materials generating force by cellular contractility^2^ and growth^3–5^, they are constantly exposed to external mechanical stresses such as tissue-extrinsic forces or boundary conditions created by neighboring tissues and structures^6^. Therefore, tissue morphology is the outcome of a complex interplay between cellular force generation, extrinsic forces, and constraints imposed by the cell’s local and global mechanical environment. Uncovering how this interplay drives the morphogenesis of growing tissues remains a challenging task.

Due to their simple structure, epithelial tissues provide an excellent model system^7^ to study how forces and material properties mechanically interact during cell shape changes that collectively determine tissue morphologies. Epithelia form sheets of tightly connected cells that are lined by a specialized extracellular matrix on their basal surface, the basement membrane (BM)^8,9^. Epithelial tissues can adopt different cellular morphologies: Squamous epithelia are composed of flat cells with a surface diameter (*D*) larger than cell height (*H*, see Figure 1A). Cuboidal epithelia consist of cube-shaped cells (*D* ≈ *H*) and columnar epithelia of cells which are higher than they are wide (*H* > *D*).

**Figure 1.**
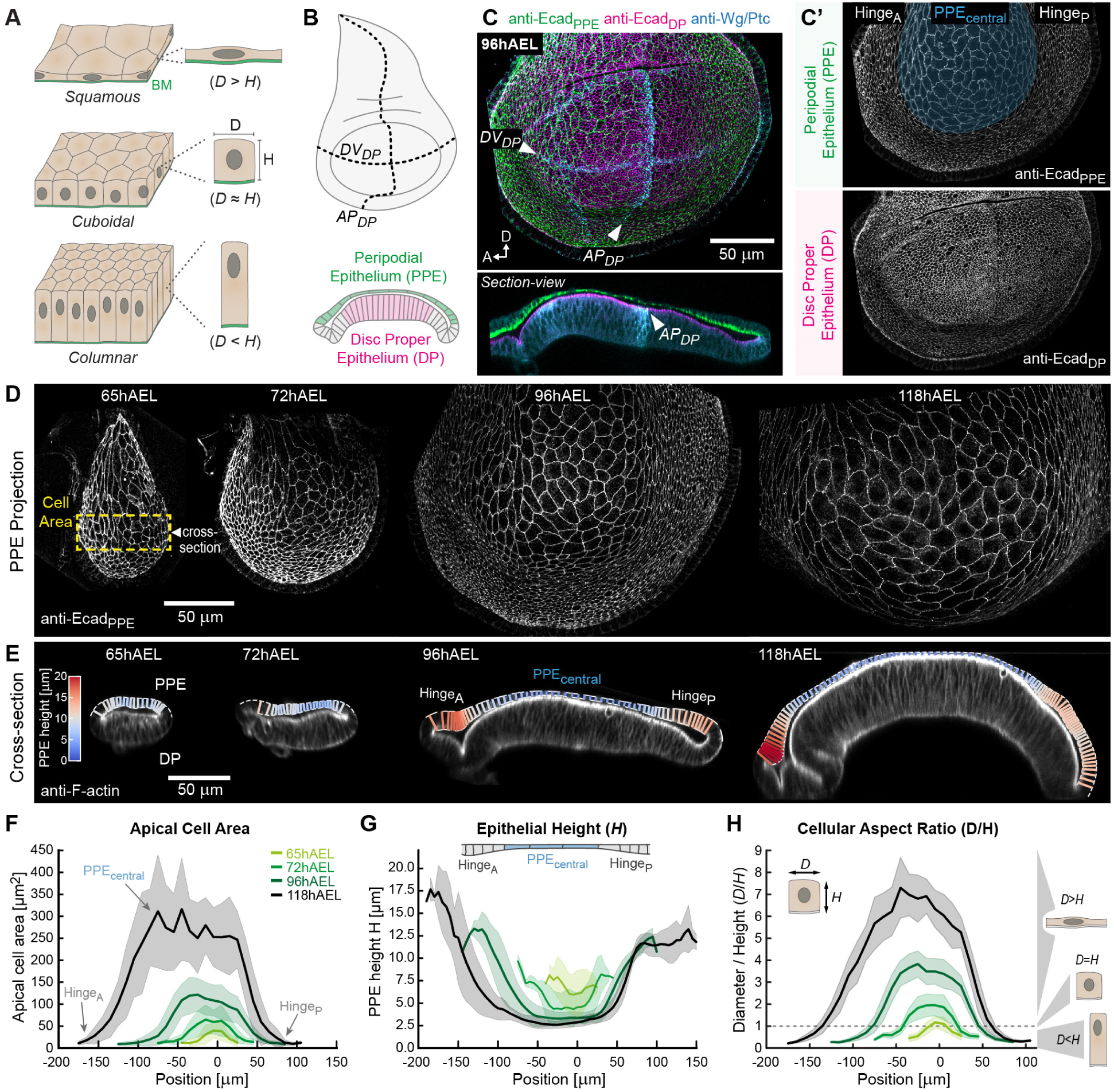
Emergence of gradients in cell morphology during peripodial morphogenesis. (**A**) Scheme showing the relation between apical cell diameter (*D*) and cell height (*H*) in the three organization types of epithelial tissues. (**B**) Scheme of the wing disc in plane (*top*) and cross-section view (*bottom*). The anterior/posterior (AP_DP_) and the dorsal/ventral (DV_DP_) compartment boundaries of the disc proper (DP) are indicated by a dashed line. (**C**) Selective-plane projections of the apical surface of the PPE (Ecad_PPE_, green) and the DP (Ecad_DP_, magenta), a cross-section view is shown below. Individual surface projections are shown in (C’). All discs are oriented with anterior (A) facing left and dorsal (D) up. In cross-sections, the PPE is facing up. (**D**) Peripodial surface projections of representative wing discs at indicated time points marked for cell outlines (Ecad_PPE_). (**E**) Cross-sections of representative wing discs with tissue outlines marked by F-actin. The PPE apical and basal surfaces are marked by dashed white lines and local tissue height is color-coded (see LUT). (**F**) Average peripodial cell area plotted along the A/P-axis (as indicated by the yellow rectangle in (D), *x*=0 corresponds to the AP_DP_). *n*-numbers [cells/discs]: *n_65_*=161/12, *n_72_*=759/12, *n_96_*=3246/13, *n_118_*=1812/12. (**G**) Plot of average epithelial height. *n*-numbers [discs]: *n_65_*=9, *n_72_*=9, *n_96_*=10, *n_118_*=8. (**H**) Plot of cellular aspect ratio (*D/H*), calculated from the data shown in (F) and (G) (see methods). *n*-numbers [cells/discs]: *n*_65_=180/12, *n*_72_=758/12, *n*_96_=3246/13, *n*_118_=1812/12. *Statistics:* Error bands in F-H indicate the standard deviation.

Epithelial tissues often transit between these shapes during development and adult life to ensure functional morphology. The cells of the urothelium (also called the transitional epithelium) are changing from cuboidal to squamous during bladder filling^10,11^. During development, cuboidal-to-squamous transitions are observed in the *Drosophila* follicular epithelium^12–14^, during *Drosophila* amnioserosa flattening^15^, and in the zebrafish retinal pigment epithelium^16^. Cuboidal-to-columnar transitions occur during the thickening of the neural epithelium^17^, during *Drosophila* wing disc bending^18,19^, and salamander neural tube invagination^20,21^. While several pathways are associated with the regulation of epithelial shape changes (including TGF-beta^12,18^, JAK/STAT^14^, JNK^22^, non-canonical dTOR^23^ and Hippo^24^ signaling), the mechanics and origin of the forces guiding these shape transitions remain largely unexplored.

Here, we use the *Drosophila* wing imaginal disc to investigate the biomechanics of epithelial cell shape transitions. The wing disc is a flattened epithelial cyst that consists of two apposed epithelial layers, the disc proper (DP) epithelium and the peripodial epithelium (PPE), connected via the peripheral hinge region (Figure 1B and 1C). This epithelial bilayer is surrounded by a BM covering the basal surface of both tissue layers. During wing disc growth, the DP epithelium increases in height, and bends to form a dome-like structure (see Fig.1C *bottom*). We previously showed that DP thickening and bending are driven by differential growth orientation between the DP epithelium which grows strictly in plane and its BM (BM_DP_), which grows more isotropic. Such differences in growth anisotropy cause geometric frustration and lead to the accumulation of compressive, elastic stress and tissue deformation^19^. Therefore, the columnar organization of the DP epithelium is induced by tissue-intrinsic, compressive forces arising from a growth orientation difference between DP tissue and its BM.

In contrast, during disc growth the central region of the overlaying PPE (PPE_central_) adopts a squamous morphology with large, flat cells and low cell density (Figure 1C’ *top*). Notably, the adjacent cells of the peripodial hinge (PPE_hinge_) do not flatten^25,26^. Input from the classical patterning signaling pathways have been associated with the squamous morphogenesis of the PPE_central_ cells^25,26^. In addition, deregulation of the putative transcription factor and cell height regulator Lines (Lin), via the mis-activation of the Dpp-target gene Spalt, transforms the PPE into a mirror image of the columnar DP^27,28^. However, more recent work points towards an important function of the peripodial BM (BM_PPE_) in the morphogenesis of the PPE layer. Mutants of the major BM component Perlecan (encoded by *trol* in *Drosophila*), exhibit increased BM stiffness and central peripodial cells fail to flatten (though discs also fail to bend)^29^. Similarly, knock-down of matrix metalloproteinase 2 (MMP2), an enzyme involved in Collagen IV cleavage and turnover, hinders peripodial flattening^19^. Overall, these recent studies point to the importance of mechanical input and particularly a role for the BM in shaping the spatially distinct tissue architecture during peripodial morphogenesis.

However, the mechanistic logic and in particular the role of the mechanical environment and the forces underlying the spatially distinct shape changes observed during peripodial morphogenesis remain uncharacterized. Here, we investigate the mechanical control of distinct epithelial cell shape transitions in the PPE during the multilayered morphogenesis of the *Drosophila* wing disc.

## Results

### Emergence of gradients in cell morphology during peripodial morphogenesis

We quantified peripodial morphology covering the major growth period of the *Drosophila* wing disc from 65 hours after egg laying (hAEL, late 2^nd^ instar) to the end of larval development (∼120hAEL at 25°C). We used the expression of Wingless (Wg), marking the DP dorsal-ventral compartment boundary (D/V_DP_), and Patched (Ptc), marking the anterior-posterior compartment boundary (A/P_DP_) as landmarks (Figure 1B+C). We quantify PPE morphology along the A/P-axis (*x*=0 corresponds to the A/P_DP_), focusing on the tissue dorsal to the D/V_DP_ (see dashed box in Figure 1D, Figure S1A and methods).

We first quantified changes in cell surface area visualized by E-Cadherin (E-cad, Figure 1D). PPE_central_ cell area increases by ∼460% from ∼50µm^2^ at 65hAEL to ∼279µm^2^ at 118hAEL (Figure 1F and Figure S1B), consistent with recent findings^24^. In contrast, the cell area of the peripheral hinge domains decreases by ∼38% (∼16µm^2^ at 72hAEL to ∼10µm^2^ at 118hAEL, Figure S1C). We next quantified peripodial height from optical cross sections (dorsal of the D/V_DP_). PPE height is initially ∼6.3µm at 65hAEL and uniform along the A/P-axis (Figure 1E and 1G) and subsequently decreases to ∼2.9µm^2^ at 118hAEL (Figure S1B). In contrast, epithelial height increases in the peripheral hinge region from ∼8µm to ∼12.9µm (Figure S1C). We next quantified cell shape changes by calculating the cellular aspect ratio (D/H, Figure 1H, using data from Figure 1F+G). Notably, the aspect ratio of PPE_central_ cells was ∼1 at 65hAEL (cuboidal shape) and subsequently increased by ∼7-fold until 118hAEL (Figure S1B), indicating a *cuboidal-to-squamous* transition. In contrast, the aspect ratio of hinge cells was ∼0.6 at 65hAEL and decreased significantly after 72hAEL, indicating a *cuboidal-to-columnar* transition (Figure S1C).

Together, these data show that peripodial morphology is the result of dynamic and spatially distinct changes in cell morphology: PPE_central_ cells undergoing a *cuboidal-to-squamous* transition while the adjacent peripheral hinge cells undergoing a *cuboidal-to-columnar* transition.

### The cuboidal-to-squamous transition is associated with an increase in cell volume

Our quantifications showed a ∼5.6-fold increase in PPE_central_ cell area between 65 and 118hAEL. Consequently, PPE_central_ cell height should decrease by ∼5.6-fold if cell volume were conserved. However, we observed that PPE_central_ height only halves (from 6.3µm to 2.9µm), suggesting that cell volume must increase by ∼3-fold. Consistently, when we estimate cellular volume by the product of cell area and height (*A*H*, data from Figure 1F and 1G) we find that PPE_central_ cell volume increases by 2.8-fold between 65 and 118hAEL (Figure 2C).

**Figure 2.**
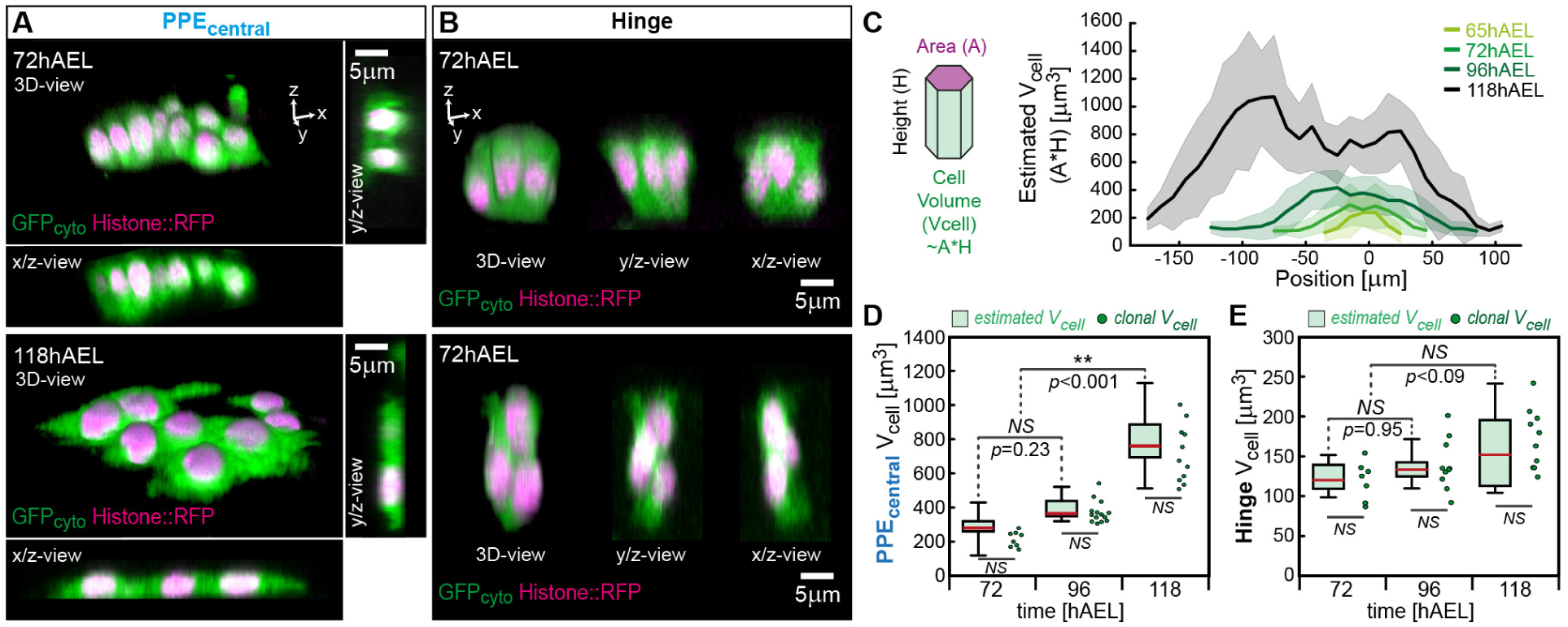
Cell volume changes during the peripodial cuboidal-to-squamous transition. (**A**+**B**) 3D rendered and cross-section views of representative PPE_central_ (A) and peripodial hinge (B) clones expressing cytosolic GFP (GFP_cyto_, green, marking the cell volume) and Histone::RFP (magenta, cell number) at 72hAEL (*top*) and at 118hAEL (*bottom*). (**C**) Peripodial cell volume (V_cell_) along the A/P-axis estimated by the product of cell area (A) and average height (H, both from Figure 1F-G). Error bands indicate standard deviation. *n*-numbers [discs]: *n_65_*=12, *n_72_*=12, *n_96_*=13, *n_118_*=12. (**D**+**E**) Average cell volume in the PPE_central_ (D) and in the peripodial hinge (E) region from volume estimations (estimated V_cell_, represented as boxplots) and from volumetric 3D segmentation of GFP-marked clones (clonal V_cell_, individual clones are plotted, see methods). *n*-numbers PPE_central_ [discs/clones]: *n_72_*=12/8, *n_96_*=13/14, *n_118_*=12/11. *n*-numbers hinge [discs/clones]: *n_72_*=7/7, *n_96_*=10/10, *n_118_*=10/10. *Statistics*: Statistical significance was assessed by a one-way ANOVA and Tukey’s post hoc test (\**p* ≤ 0.05, \*\**p* ≤ 0.005, \*\*\**p* ≤ 0.0005). In box plots, the median is indicated by a central thick line, while a box outlines the interquartile range (containing 50% of the data points). Whiskers indicate the minimum and maximum data range.

We therefore quantified cellular volume by expressing cytosolic GFP (GFP_cyto_) in sparse clones of cells (Figure S1D). Notably, clonal labeling well recapitulates the expected cell shape changes (Figure 2A-B). We quantified average cell volumes in the PPE_central_ (Figure 1D) and the hinge region (Figure 1E) by 3D volume segmentation of our clonal data (clonal V_Cell_) and compared it to the estimated cell volumes (estimated V_cell_). Both approaches consistently showed that the volume of PPE_central_ cells increased from ∼290µm^3^ at 72hAEL to ∼800µm^3^ at 118hAEL (Figure 2D). In contrast, hinge cell volume did not change significantly (Figure 2E).

These results indicate inverse gradients in cell density (low central to high peripheral) and volume (high central to low peripheral) raising two major questions: (1) What regulates the inverse gradients in cell density and volume? (2) What physical forces are responsible for the inverse shape transitions of PPE_central_ versus hinge cells?

### Endoreplication of squamous cells controls cell volume and cell shape

We first addressed the cause of the cell density gradient and its relation to the increase in PPE_central_ cell volume. We hypothesized that the cell density gradient (low central to high peripheral) could stem from a non-uniform proliferation pattern. We quantified proliferation rates using the mitotic marker Phospho-Histone-H3 (PHH3, Figure 3A). The mitotic rate was higher in central cells than in hinge cells at 65hAEL. Subsequently, the mitotic rate of central cells decreased after 72hAEL and dropped to zero at 118hAEL (Figure 3B, dark and light blue, see methods). In contrast, hinge cells maintained a positive mitotic rate throughout disc growth and continued to proliferate even at 118hAEL (Figure 3B, gray).

**Figure 3.**
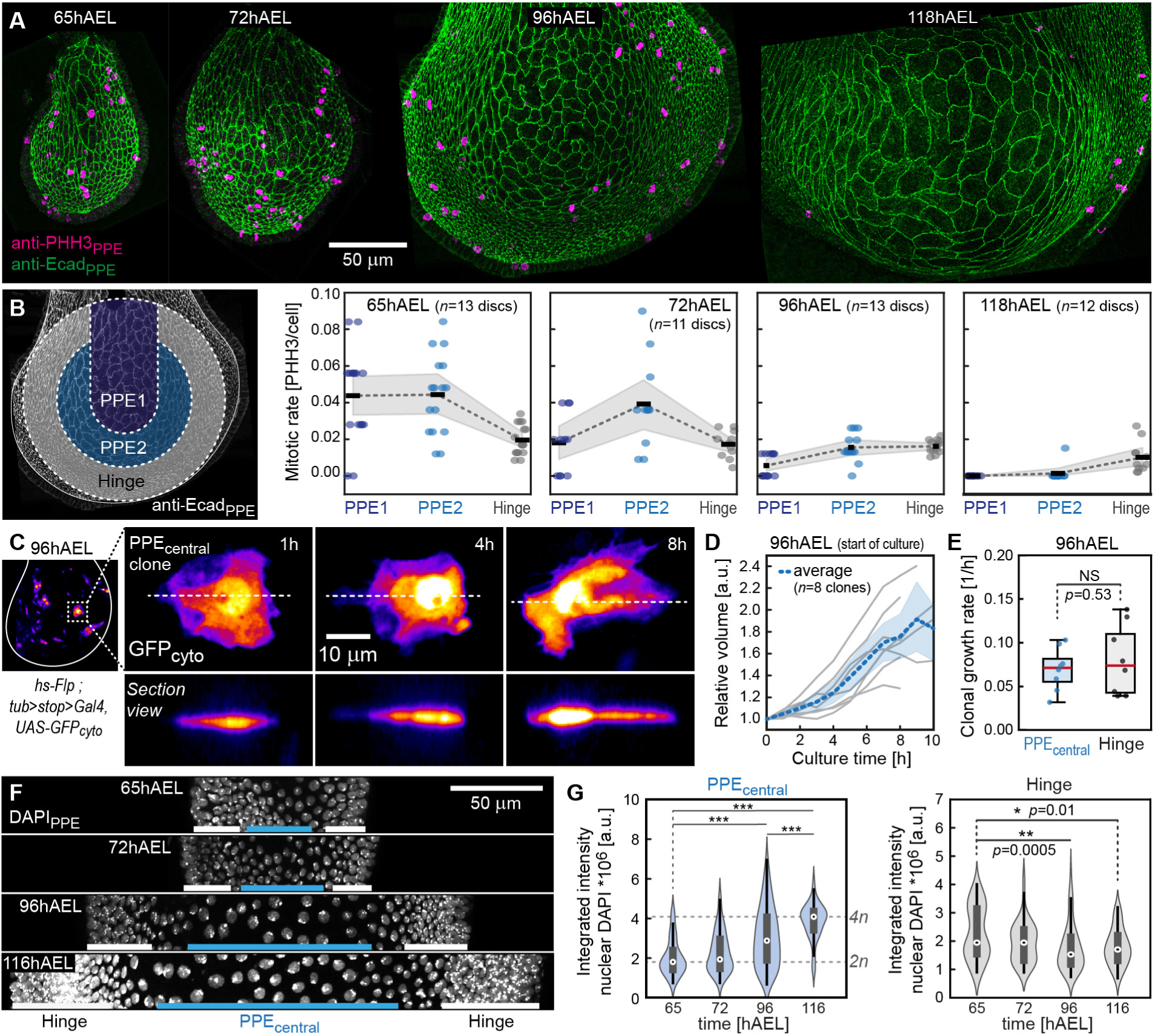
Endoreplication of squamous cells controls cell volume and cell shape. (**A**) Representative peripodial projections of wing disc stained for PHH3 (magenta) and Ecad (green). (**B**) Quantification of mitotic rate (PHH3/cell, see methods) in three circular domains. PPE1 (dark blue) and PPE2 (light blue) correspond to the squamous PPE_central_ while the peripheral ring (gray) covers the peripodial hinge domain. (**C**) *left*: 96hAEL old wing disc expressing cytosolic GFP (*GFP_cyto_*) in a clonal manner at the beginning of *ex vivo* culture. *right*: A representative PPE_central_ clone is shown in projection (*top*) and cross-section view (*bottom*) at indicated times of *ex vivo* imaging. (**D**) Quantification of the relative volume change of 8 PPE_central_ clones during *ex vivo* culture. Individual clonal growth curves are shown in grey and the average growth curve in blue (error band indicates standard deviation). (**E**) Average clonal growth rate in the PPE_central_ (blue) and hinge region (grey). *n*-numbers [clones/discs]: *n*_central_=8/2, *n*_hinge_=8/2. (**F**) Projections of the PPE tissue overlaying the D/V_DP_ of discs stained for DAPI (grey). PPE_Central_ and hinge cells are indicated by blue and white boxes, respectively. (**G**) Temporal changes of the integrated nuclear DAPI intensity (∼DNA content) obtained from 3D nuclear volume segmentation for the PPE_central_ (*left*) and the hinge (*right*, see methods). In violin plots, a spot indicates the median, a thick grey line the interquartile range, and a black line the minimum and maximum data range. Data distribution is visualized by a kernel density estimation. *n*-numbers [disc/nuclei] for PPE_central_: *n_65_*=5/79, *n_72_*=5/79, *n_96_*=7/122, *n_116_*=8/103 and for the hinge: *n_65_*=5/63, *n_72_*=5/49, *n_96_*=7/91, *n_116_*=5/46. *Statistics*: Statistical significance was assessed by a one-way ANOVA and Tukey’s post hoc test ( \**p* ≤ 0.05, \*\**p* ≤ 0.005, \*\*\**p* ≤ 0.0005).

Therefore, PPE_central_ cells stop to divide around 96hAEL, corresponding to the time at which PPE_central_ cell volume starts to increase (Figure 2D) and central cell density to decrease (Figure S1B). Hence, continued cellular growth must drive the subsequent increase in PPE_central_ cell volume. To this end, we quantified the cellular volume changes of PPE_central_ cells marked by clonal expression of cytosolic GFP from live imaging movies of *ex vivo* cultured discs (Figure 3C, see methods). We quantified clone volume in up to 10h long movies and consistently observed increasing clone volume (Figure 3D) and a positive growth rate (Figure 3E) after 96hAEL. Therefore, squamous cells continue to grow in volume despite exiting the cell cycle. Notably, the growth rate of PPE_central_ cells did not differ from peripheral hinge cells (Figure 3E), indicating that not differential growth but differential cell division dynamics drive the emergence of a cell density gradient during PPE morphogenesis.

Cellular volume within cell types is known to be tightly regulated^30^ and directly related to the cell’s DNA content^31^. Excessive cell growth leads to cytoplasmic dilution negatively impacting cell homeostasis and proliferation^32^. A commonly employed process that allows cells to increase their volume is to increase the cell’s DNA content via endoreplication^31,33^. To ask if squamous cells undergo endoreplication we assessed their DNA content by quantifying the nuclear-integrated DAPI intensity (Figure 3F-G, see methods). We observed that the integrated DAPI intensity in squamous cells ∼doubled from 65 to 118hAEL (Figure 3G, left), suggesting that squamous cells increase their nuclear DNA content from a ploidy value of 2C to 4C by endoreplication^33^. In contrast, peripheral hinge cells maintain a diploid cell fraction throughout development (Figure 3G, right).

This shows that the peripodial cell density gradient stems from a spatially non-uniform cell division pattern, rather than non-uniform growth rates, allowing squamous cells to increase their volume due to continued growth enabled by endoreplication.

### Wing disc bending stretches the squamous cells leading to cell flattening

Next, we investigated the forces guiding the *cuboidal-to-squamous* transition of PPE_central_ cells. During disc growth, the initially flat disc bends to form a dome-like structure towards the end of development^19^ (Figure 4A). Simple approximations suggest that doming after 72hAEL leads to a greater expansion of the surface area that the PPE covers than expected by the volume increase of the PPE layer (Figure S2B). Hence, the peripodial area expands faster than it can compensate for by growth. We therefore hypothesized that the doming DP exerts a bending stress on the PPE layer leading to its elastic deformation and cell flattening (Figure 4A, *middle*).

**Figure 4.**
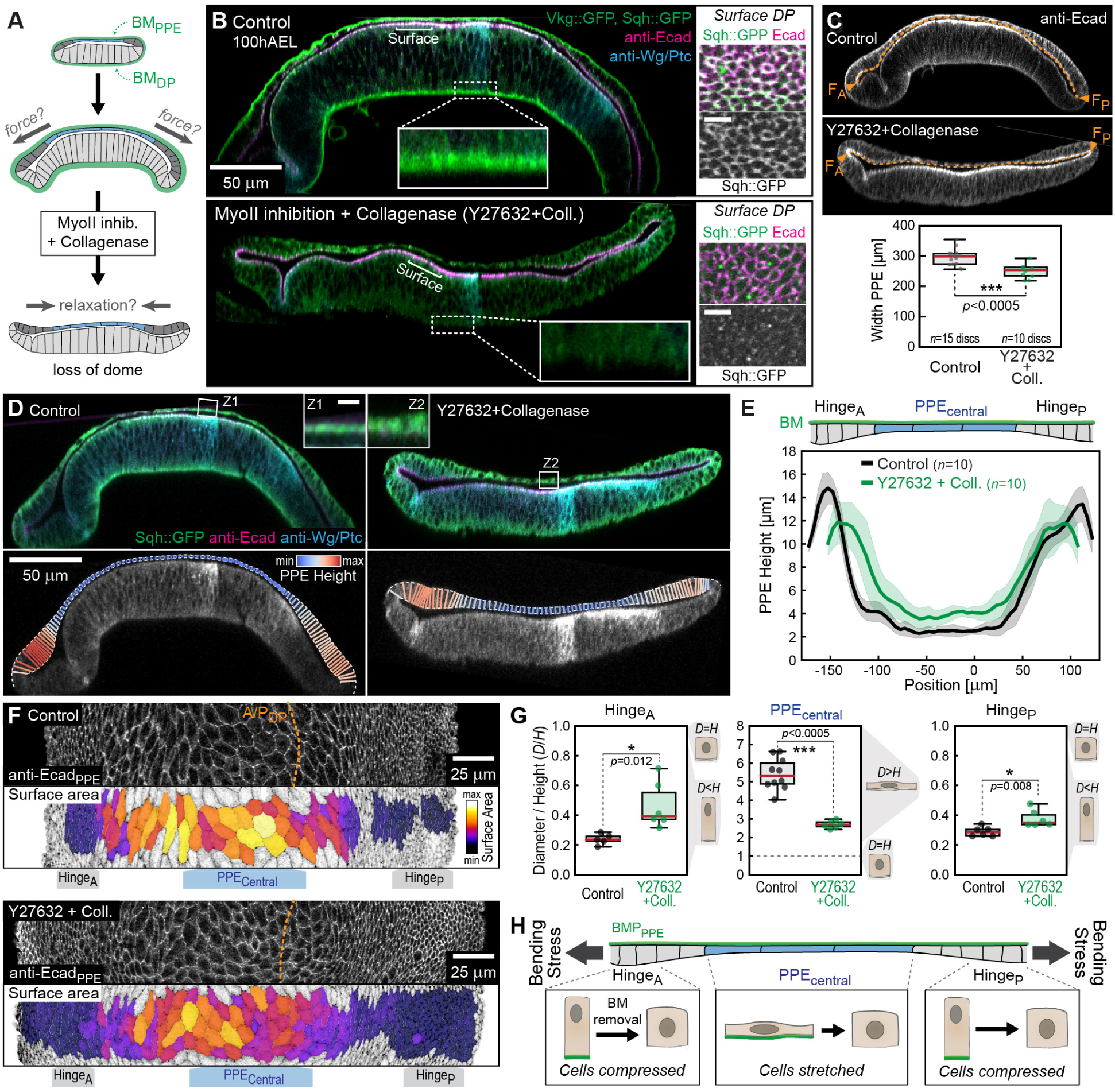
DP bending stretches the squamous cells leading to cell flattening. (**A**) The initially flat wing disc (*top*) bends during its growth phase, leading to stretching of the PPE layer (*middle*). MyoII inhibition and BM digestion by Collagenase result in tissue relaxation and flattening (*bottom*). (**B**) Sections of representative wing discs marked for Vkg::GFP, MyoII (Sqh::GFP, both green) and cell outlines (Ecad, magenta): non-treated (control, *top*) and treated with Rock inhibitor (Y27632) and Collagenase (*bottom*). DP surface projections at the indicated position are shown on the right (scale bar = 5µm). (**C**) Cross-sections of control (*top*) and Y27632 + Collagen-treated discs (*middle*). The width of the PPE apical surface between the anterior (F_A_) to the posterior fold (F_P_) is indicated by dashed orange lines and quantified at the bottom. (*p*=0.00021). (**D**) Cross-sections of representative 96hAEL control (*left*) and Y27632+Coll. discs (*right*), inserts show magnifications of the PPE layer above the AP_DP_ (scale bar = 4µm). Local PPE height is color-coded (*bottom*). (**E**) Quantification of PPE height along the A/P-axis. Error bands indicate standard deviation. (**F**) PPE surface projection of control (*top*) and Y27632+Coll. discs (*bottom*) labelled for cell outlines (Ecad_PPE_). Overlays of cell outlines (black) with quantified cell surface area (color-coded) are shown (*lower*). Approximate domains of Hinge_A_, PPE_central_ and Hinge_P_ are indicated. (**G**) Quantification of cell aspect ratio (D/H). *n*-numbers [disc] for Hinge_A_: *n_cont._*=5, *n_Y+C_*=6; PPE_central_: *n_cont._*=10, *n_Y+C_*=6; Hinge_P_: *n_cont._*=6, *n_Y+C_*=6. (**H**) Stress distribution in the PPE tissue layer: The PPE layer is stretched by disc bending leading to tensile stress in the PPE_central_ cells, while cells in the hinge region are compressed. *Statistics*: In box plots, the median is indicated by a central thick line, the interquartile range (50% of data points) by a box and data range by whiskers. Statistical significance was assessed by a two-sided Student’s *t*-test (unequal variance, \**p* ≤ 0.05, \*\**p* ≤ 0.005, \*\*\**p* ≤ 0.0005).

To test this hypothesis, we reverted disc bending and evaluated changes in peripodial morphology upon elimination of the bending-induced stress. Notably, disc bending crucially depends on the BM of the DP layer (BM_DP_), visualized by a GFP-tagged version of *Drosophila* Collagen IV, called Viking (Vkg::GFP, see Figure 4B, *top*). The acute digestion of the BM by Collagenase and simultaneous inhibition of Myosin II contractility using the Rock inhibitor Y27632 (referred to as Y+C discs, see methods) leads to a loss of bending and disc flattening (Figure 4B *bottom*, also see ^19,34^). Upon tissue flattening the width of the PPE layer decreased by ∼15.5% (Figure 4C) confirming that the PPE is stretched elastically by a tissue-extrinsic force originating from DP bending.

We next asked how the bending stress affects the different domains of the PPE (PPE_central_ and the anterior and posterior parts of the hinge, or Hinge_A_ and Hinge_P_). We quantified changes in PPE height along the A/P-axis in control (domed and stressed) and Y+C discs (flat and relaxed). The height of PPE_central_ cells increased by 60% upon disc flattening from ∼2.5 in control to ∼3.9µm in Y+C discs (Figure 4D-E and Figure S3A-C). This indicates that squamous PPE_central_ cells are elastically stretched by a planar tensile stress originating from DP bending. Notably, this relaxation behavior is absent at 65hAEL but is observed from 72hAEL onwards (Figure S3D-F), corresponding to the time point when central PPE cells start to undergo a cuboidal-to-squamous transition (see Figure 1H). In contrast, we observed an inverse relaxation behavior in the hinge regions. Epithelial height decreases in both the Hinge_A_ and Hinge_P_ by 18% and 16%, respectively. This indicates the release of compressive stress in the hinge tissue upon removal of the BM (Figure 4D-E and Figure S3B-C). Notably, the observed changes in tissue morphology upon disc flattening are consistent with the observed changes in cell shape (Figure 4F-G).

In summary, these results indicate that a tensile stress due to disc bending leads to elastic stretching and flattening of the squamous PPE_central_ cells. In contrast however, the hinge tissue is compressed, indicating that hinge cells are not stretched by DP bending (Figure 4H). This raises the question of why the response to the external bending force differs between the central squamous cells and the adjacent peripheral hinge regions.

### Mechanical differences in the BM underly central squamous flattening versus hinge compaction

We first considered the possibility that the BM mechanically protects hinge cells from the bending stress. We previously showed that anisotropies in BM_DP_ expansion relative to DP growth lead to geometric frustration, compressive stress and cell compaction in the DP^19^. Analogously, we hypothesized that the hinge BM_PPE_ could oppose planar hinge expansion and growth, leading to compressive stress and cell compaction in the hinge.

Accordingly, the growing hinge cells exert a tensile stress on the underlying hinge BM_PPE_, which is further amplified by the stress due to DP bending. To assess tensile stress in the BM_PPE_ we eliminated the forces exerted by planar hinge growth and DP bending by the acute removal of both cell layers. The cell layers were removed by *ex vivo* exposure of discs to the detergent Triton X-100 (referred to as decellularization, see methods and ^19^), leading to lipid bilayer degradation and the loss of cellular stresses acting on the BM_PPE_ (Figure 5A-B). To follow BM_PPE_ relaxation we photo-bleached circles on the Vkg::GFP_PPE_ labeling. Notably, changes in circle area indicate stress relaxation in the BM: A reduction in circle area upon decellularization indicates that the BM was under tensile stress, while no change in area indicates that the BM was stressless, i.e. relaxed (Figure 5C, *left*). We bleached circles in the hinge_A_, PPE_central_ and hinge_P_ regions of 96hAEL wing discs and observed changes in circle area before (A_0_) and 60 min after decellularization (A_60_, see Figure 5B-C). We did not observe changes in circle area in the central BM_PPE_, indicating that the central portion of the peripodial BM_PPE_ was not stressed (in contrast to the tensile stress in the PPE_central_ epithelial layer, see above). However, following decellularization, circle area decreased notably in the BM_PPE_ of both the hinge_A_ and hinge_P_ region by 22% and 9%, respectively (Figure 5C’). Furthermore, the extent of BM_PPE_ relaxation positively correlated with the initial epithelial height (Figure 5C’’). These experiments show that the hinge BM_PPE_ is stretched by the growing hinge epithelium, raising the question why the hinge BM_PPE_ accumulates elastic stress, while the central BM_PPE_ does not. Previously we showed that increased BM thickness is associated with accumulation of tensile stress in the BM_DP_ of the DP epithelium^19^. Indeed, we found that the BM_PPE_ of hinge cells is thicker than the BM_PPE_ of squamous cells (Figure 5D). Hence, like in the DP, a growth differential between the hinge cells and the associated BM_PPE_ drives cell compression and an increase in hinge cell height (Figure 5E).

**Figure 5.**
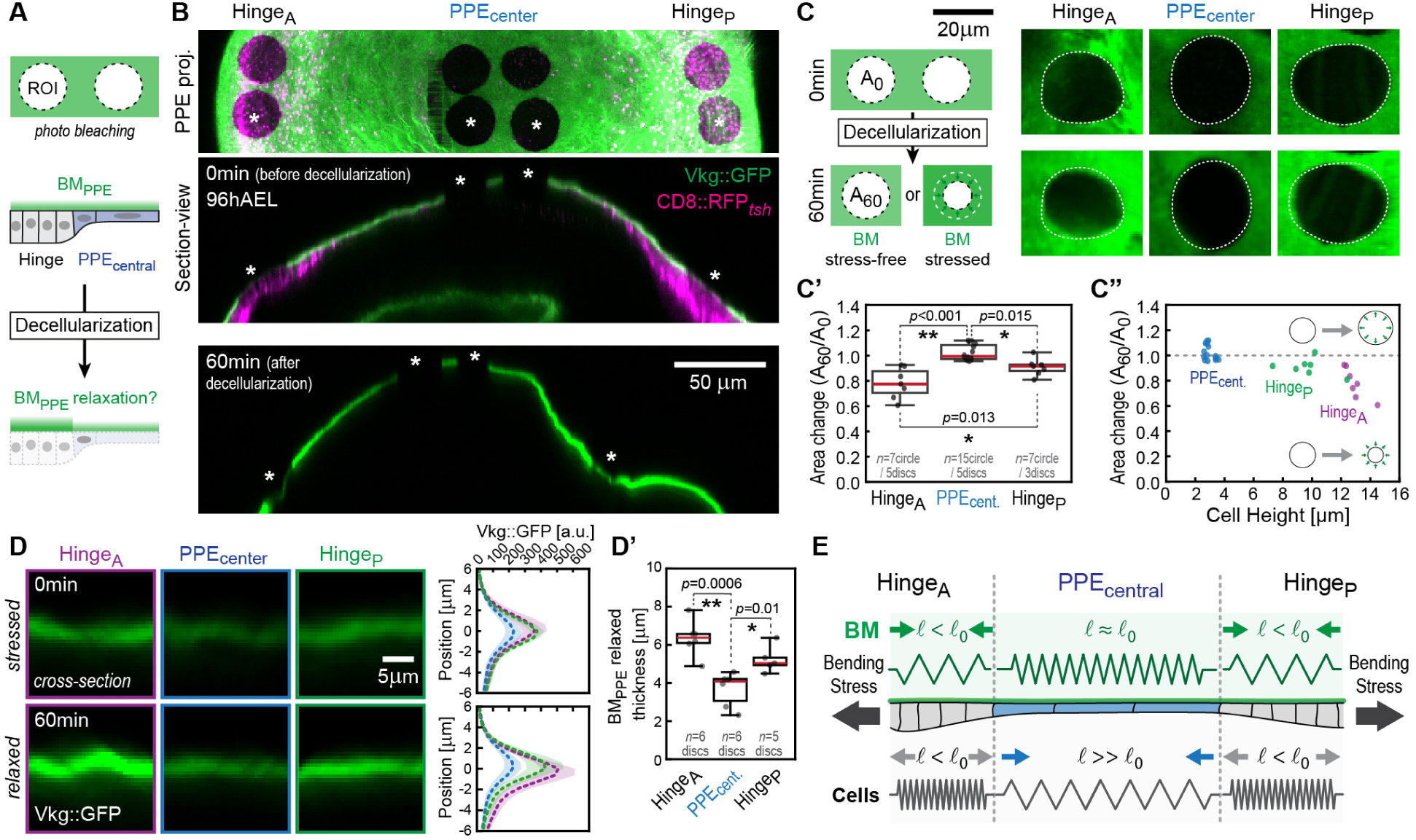
– Mechanical differences in the BM cause central squamous flattening versus hinge compaction. (**A**) Scheme of the BM_PPE_ (green) in surface (*top*) and section-view (*middle-bottom*) and underlying cells (hinge grey, PPE_central_ blue). Circular regions of interest (ROI) are photo-bleached on the Vkg::GFP signal, allowing to follow BM_PPE_ relaxation upon decellularization (*bottom*). (**B**) Projection (*top*) and section views of a 96hAEL Vkg::GFP wing disc expressing RFP in the PPE layer (*tsh-Gal4*) before (*top, middle*) and 60min after decellularization (*bottom*). Bleached ROIs are marked by asterisks. (**C**) *left*: The initial circle area (A_0_) either decreases or remains unchanged upon decellularization (A_60_) depending on if the BM_PPE_ was stressed or relaxed, respectively. *right*: Bleached circles in indicated regions before (0min) and after decellularization (60min). Original circle area is indicated by dashed line. (**C’**). Quantification of relative circle area 60min after decellularization. (**C’’**) Relative circle area 60min after decellularization plotted against the initial epithelial height. (**D**) Cross-sections of the BM_PPE_ (Vkg::GFP) in indicated regions before (*top*, stressed) or after decellularization (*bottom*, relaxed). Average intensity profiles are shown to the right. (**D’**) Quantification of relaxed BM_PPE_ thickness (see methods). (**E**) Scheme of the stresses present in the distinct structures of the PPE due to external bending stress and internal growth-induced stress due to hinge expansion. Stresses are illustrated by arrows and for conceptual support visualized by springs, where l_0_ corresponds to the relaxed rest length of the spring. Therefore, l < l_0_ corresponds to compressive stress and l > l_0_ to tensile stress. *Statistics*: In (D) error bands indicate the standard deviation. In box plots (C’+D’) the median is indicated by a central thick line, the interquartile range (50% of data points) by a box, and data range by whiskers. Statistical significance was assessed by a one-way ANOVA and Tukey’s post hoc test (\**p* ≤ 0.05, \*\**p* ≤ 0.005, \*\*\**p* ≤ 0.0005).

We next investigated if the hinge BM_PPE_ is additionally stressed by DP bending. We genetically digested the BM_DP_ by overexpression of Matrix Metalloproteinase 2 (MMP2), an enzyme that degrades Collagen^35–37^ specifically in the DP. MMP overexpression in the DP for 7h (using the Gal80 temperature-sensitive system^38^, see methods) and simultaneous inhibition of MyoII (by Y27632) resulted in flattened wing discs that had lost the BM_DP_ but maintained the BM_PPE_ (referred to as MMP2_OE_ + Y276932, see Figure S4). Notably, Vkg::GFP_PPE_ mean intensity doubled in the hinge regions upon disc flattening (see asterisks in Figure S4F) indicating an increased density of the hinge BM_PPE_ and the presence of a tensile stress in the hinge BM_PPE_ that relaxes elastically upon disc flattening.

In summary, we show that the hinge BM_PPE_ is mechanically stressed by (i) an external stress from disc bending and by (ii) a growth-induced stress from hinge expansion (Figure 5E). Therefore, the *cuboidal-to-columnar* transition in the hinge stems from the BM_PPE_ that shields hinge cells from the bending stress and from hinge cells growth-induced compressive stress. Notably, we find that the mechanical state and thickness of the BM_PPE_ differs distinctly between the PPE_central_ and the hinge domain: While the hinge BM_PPE_ is thick and stressed, the central BM_PPE_ is thin and stressless and thereby does not shield the overlying PPE_central_ cell layer from external bending stresses, leading to cell flattening (see discussion).

### Mechanical coupling mediated by Dumpy leads to shearing of the squamous cells

These data emphasize the role of inter-tissue stress propagation. Thus, we next addressed how the bending stress mechanically propagates from the bending DP to the PPE layer. We showed that DP bending leads to tensile stress in the hinge BM_PPE_ (Figure S4), supporting the transmission of tensile stress via the hinge region linking the DP and the PPE at their periphery (Mechanism I, Figure 6A, *top*). In addition, the apical surfaces of the DP and the PPE appear in direct physical contact possibly leading to the shearing of the PPE by the bending DP layer (Mechanism II, Figure 6A, *bottom*). Despite their physical proximity, it remains unknown if a direct mechanical connection exists between the two layers.

**Figure 6.**
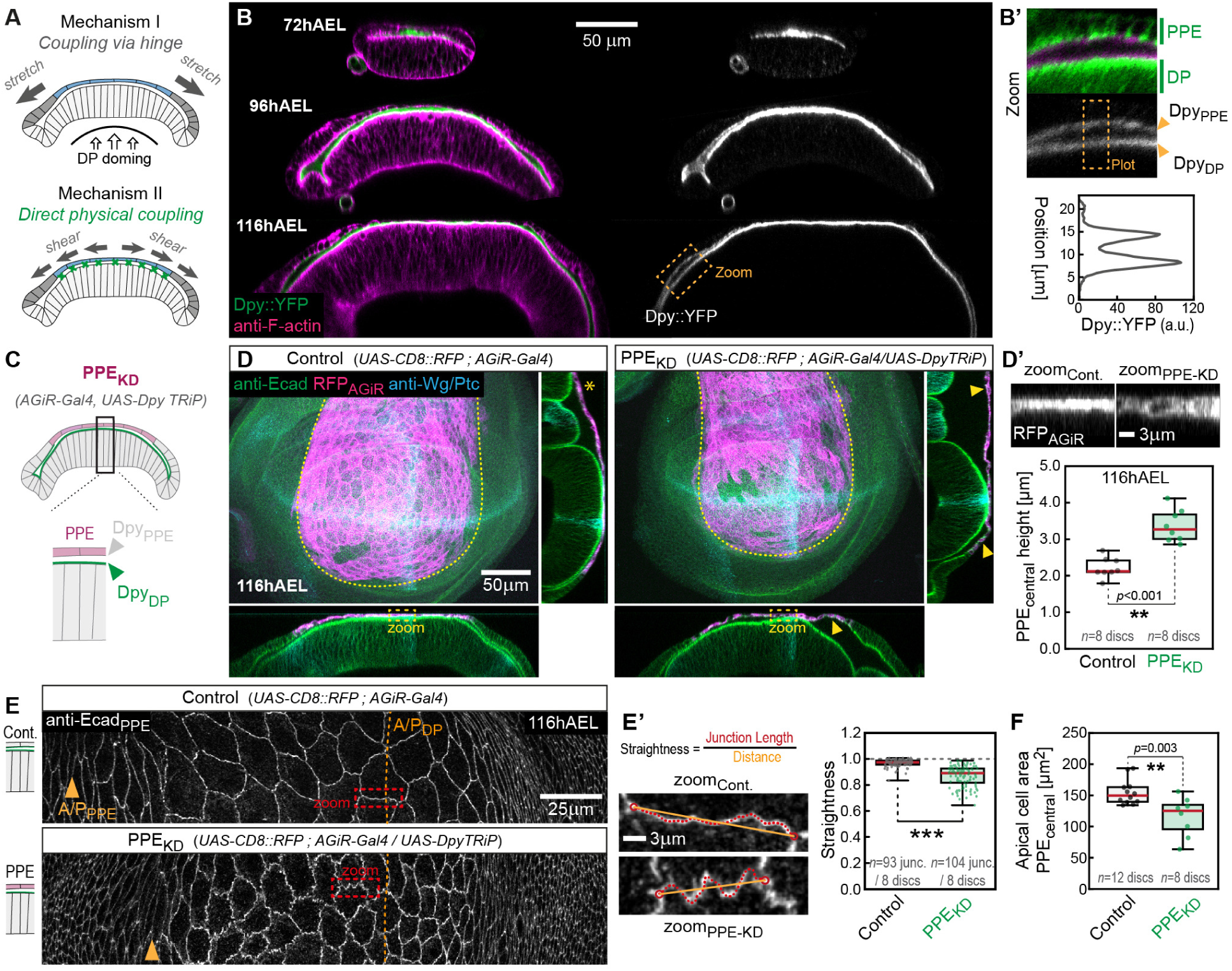
Mechanical coupling mediated by Dpy leads to shearing of squamous cells. (**A**) Stress due to DP bending is transmitted via the hinge region (*Scenario I*) and/or by direct physical coupling, shearing the PPE (*Scenario II*). (**B**) Sections of Dpy::YFP (green) wing discs stained for F-Actin (magenta). In the zoom (B’) orange arrowheads indicate Dpy localizing along the surface of the PPE (Dpy_PPE_) and the DP (Dpy_DP_). Intensity is plotted below. (**C**) Knock-down of Dpy in the PPE_central_ cells (by *AGiR-Gal4*) leads to a loss of Dpy_PPE_. (**D**) Maximum projection (*centre*) and *x-z* and *y-z* section-views (*bottom* and *right*, respectively) of wing discs expressing RFP in the central PPE cells (magenta), either alone (control, *left*) or together with a DpyTRiP line (PPE_KD_, *right*). Arrowheads mark sections where the PPE has detached from the DP. PPE_central_ height is visualized and quantified in (D’). (**E**) Cell outlines marked by anti-Ecad (white) in peripodial projections of a control (*top*) and a PPE_KD_ disc (*bottom*). Orange arrowheads mark the A/P_PPE_. The junctional regions marked by dashed boxes are enlarged in (E’). Junctional straightness is quantified on the right. (**F**) Quantification of PPE_central_ cell area. *Statistics*: Red line in box plots indicates the median, boxes mark the interquartile range (50% of data points) and whiskers the data range. Statistical significance was assessed by a two-sided Student’s *t*-test (unequal variance, *P ≤ 0.05, **P ≤ 0.005, ***P ≤ 0.0005).

The apical extracellular matrix protein Dumpy^39^ (Dpy) is a good candidate for connecting the two layers. Dpy is known to link epithelial^40,41^ and tendon cells^42^ to the cuticle and to align bristle orientation^43^ during metamorphosis and contains a zona pellucida domain^44^, capable of forming disulfide-linked homodimers^45^ or filaments^42^. While Dpy is expressed in the early pupal wing disc^37,46^, Dpy expression and function was not investigated during larval development. Using a Dpy protein-trap line (Dpy::YFP)^47,48^ we observed that Dpy::YFP is already expressed at 72hAEL (Figure 6B and Figure S5A). Notably, Dpy is a transmembrane protein and localizes to the apical surface of both, the DP (Dpy::YFP_DP_) and the PPE (Dpy::YFP_PPE_, see Figure 6B’).

To study Dpy function during peripodial morphogenesis, we knocked down Dpy specifically in the squamous PPE_central_ cells using *AGiR-Gal4* (referred to as PPE_KD_, Figure 6C-D and Figure S5B-C). While the apical surface of the PPE and the DP layer are usually in close contact, in PPE_KD_ disc a larger luminal space separated the two epithelial layers (see arrowhead in Figure 6D and Figure S6A+C). Furthermore, the cuboidal-to-squamous transition of PPE_central_ cells was impaired in PPE_KD_ discs, indicated by an increase in epithelial height from 2.4µm to 3.7µm (Figure 6D’ and Figure S6D), relaxed squamous cell junctions (Figure 6E) and reduced apical surface area (Figure 6F). These observations indicate that Dpy physically connects the DP and PPE, and that this may lead to the transmission of shear stress from the doming DP to PPE_central_ cells and cell flatterning. Consistently, we observed that the morphology of the anterior hinge region is altered in PPE_KD_ disc (Figure S6E), consistent with ‘slippage’ between the two epithelial layers due to impaired coupling and increased tensile force transmission via the hinge.

In summary, these results show that the elastic flattening of squamous PPE_central_ cells depends on shear-stress originating from DP bending that is transmitted by direct physical coupling via the apical extracellular matrix protein Dpy.

### Dpy is required to tune the mechanical state of the peripodial BM

Loss of Dpy from the apical surface of the PPE layer impairs epithelial linkage and shear force transmission to the PPE. However, as Dpy is also expressed in DP cells we next addressed the role of Dpy localization to the apical surface of the DP. We therefore knocked down Dpy in the posterior compartment of the wing disc using *hedgehog-Gal4* (*hh-Gal4*), referred to as Posterior_KD_ (Figure S5B-C).

In posterior_KD_ discs Dpy::YFP was absent in the posterior compartment from the apical surface of both epithelial layers (Figure S5C, *bottom*). Notably, the A/P-compartment boundary in the PPE layer (A/P_PPE_) is shifted anteriorly with respect to the DP epithelium (A/P_DP_, see Figure 7A *left*). Therefore, Dpy is lost from both epithelial layers (DP and PPE) only in the tissue posterior (right) of the A/P_DP_, which we refer to as the Dpy_DP-/PPE-_ domain. We observed a significant increase in PPE height restricted to the Dpy_DP-/PPE-_ domain (Figure 7A *right*). This height increase was accompanied by an increase in peripodial cell density (Figure 7A’) and a transition in cellular aspect ratio to columnar cell shape (Figure 7B), completely blocking squamous morphogenesis. Furthermore, we observed frequent cell division events in the Dpy_DP-/PPE-_ domain even at 116hAEL (not shown), while control discs stop dividing after 96hAEL (Figure 3A-B). These observations hinted to alterations in signaling activity in the Dpy_DP-/PPE-_ domain. Ectopic Decapentaplegic (Dpp) signaling is known to induce proliferation^49,50^ and increase cell height^18,28^. Indeed, we found that Dpp signaling, visualized by phosphorylated Mad (P-Mad), is significantly upregulated in the Dpy_DP-/PPE-_ domain (Figure S7 and Figure S8A-C). However, peripodial overexpression of a constitutive active form of the Dpp Type-I receptor Thickveins (TkvCA) only caused a small (25%) increase in tissue height (from 2.3 to 2.9µm, Figure S8E-F), indicating that additional signaling inputs drive the 6.5-fold cell height increase in Posterior_KD_ discs (from 2.3 to 15µm, Figure S6D’). We finally investigated how the loss of Dpy in posterior_KD_ discs blocks the cuboidal-to-squamous transition of PPE_central_ cells. We showed that squamous morphogenesis requires a stressless BM_PPE_ that relays external tensile stresses onto the underlying PPE cell layer. We therefore investigated BM structure and mechanics in posterior_KD_ discs using Vkg::GFP. The Vkg::GFP_PPE_ intensity was increased in the Dpy_DP-/PPE-_ domain (Figure 7C), consistent with changes in BM structure and mechanics. To assess changes in BM_PPE_ stress levels in posterior_KD_ discs we bleached circles anterior (control, low cell height) and posterior of the A/P_DP_ (increased cell height) and subsequently decellularized the discs and followed BM_PPE_ relaxation (Figure 7D-F). Upon decellularization anterior control circles did not change in area (Figure 7E), confirming that the BM_PPE_ of squamous cells is not stressed. In contrast, in the Dpy_DP-/PPE-_ domain (where cell height is increased) circle area decreased significantly (Figure 7E), indicating the presence of tensile stress in the BM_PPE_ covering the Dpy_DP-/PPE-_ domain. Consistently, we find that the BM_PPE_ thickness in the Dpy_DP-/PPE-_ domain was increased (Figure 7F).

**Figure 7.**
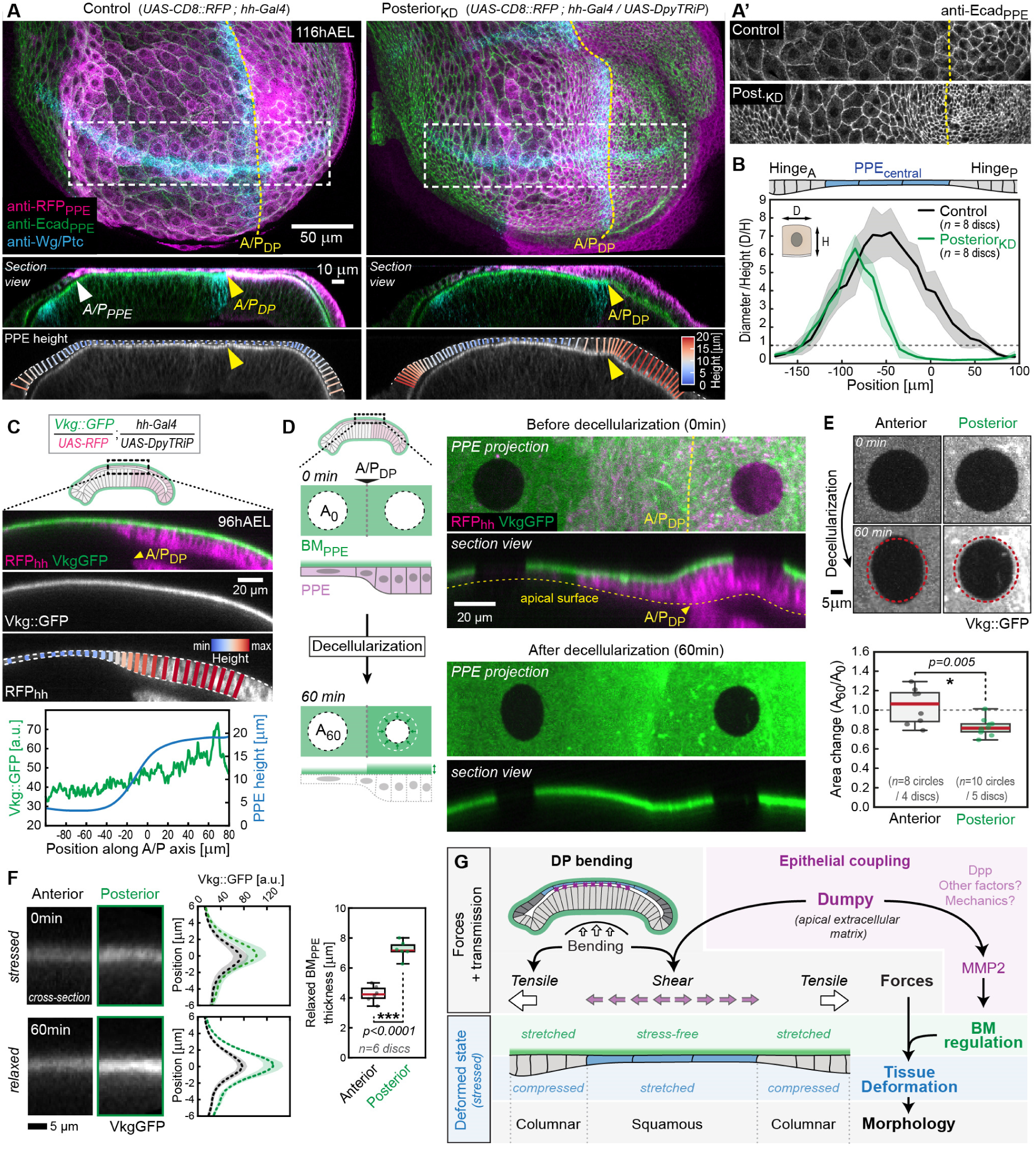
Dpy is required to tune the mechanical state of the peripodial BM. (**A**) PPE projections (*top*) and section views (*bottom*) of discs expressing RFP (Control, *left*) or RFP and Dpy-TRiP (Posterior_KD_, *right*) in the posterior compartment. PPE height is colour coded. Regions marked by white rectangles are magnified in (A’). (**B**) Plot of aspect ratio (D/H) of the genotypes shown in (A). (**C**) Cross-section view of a Posterior_KD_ disc (*top*). The BM_PPE_ is marked by Vkg::GFP (green) and PPE height is colour coded. *Bottom*: Quantification of Vkg::GFP intensity (green) and PPE height (blue). (**D**) *left*: Scheme of circular bleaching and decellularization in Posterior_KD_ discs anterior to the A/P_DP_ (low cell height) and posterior to the A/P_DP_ (increased cell height). Upon decellularization, the circular area will decrease (A_60_<A_0_) while BM thickness and intensity will increase if the BM was under tensile stress. *right*: PPE projection and section view of a Posterior_KD_ wing disc expressing Vkg::GFP before (0min, *top*) and after decellularization (60 min, *bottom*). The A/P_DP_ is marked by a yellow dashed line (PPE projection) and an arrowhead (cross-section). (**E**) Representative circles before (0min) and after decellularization (60min). Relative circle area upon decellularization is quantified below. (**F**) Cross-section views of the BM_PPE_ (Vkg::GFP) at indicated positions before (*top*) and after decellularization (*bottom*) and quantification of Vkg::GFP intensity (*middle*) and relaxed BM_PPE_ thickness (*left*). (**G**) Schematic summary: Cell shape changes result from deformations driven by the interplay of bending force transmission (tensile and shear) and the spatial regulation of BM_PPE_ mechanics. *Statistics*: Error bands in (B) indicate standard deviation. The red line in box plots in (E-F) indicates the median, boxes mark the interquartile range (50% of data points), and whiskers the data range. Statistical significance was assessed by a two-sided Student’s *t*-test (unequal variance, *P ≤ 0.05, **P ≤ 0.005, ***P ≤ 0.0005).

We previously showed that MMP2 expression in the PPE is essential to drive planar, stressless BM_PPE_ growth, whereas a loss of MMP2 leads to increased BM_PPE_ thickness, the accumulation of compressive stress and cell height increase^19^. Consistently, we found that in posterior_KD_ discs MMP2 levels were significantly reduced in the Dpy_DP-/PPE-_ domain (Figure S9A-D). Notably, peripodial MMP2 levels negatively correlated with epithelial height (Figure S9C). Thus, Dpy is required in at least one layer (DP or PPE) for peripodial MMP2 expression which tunes stress-free BM_PPE_ expansion and squamous morphogenesis.

In summary, we here showed that Dpy, in addition to coupling epithelial layers, is required to establish signaling crosstalk between the two epithelial layers to control BM mechanics and thereby to allow expansion and flattening of squamous cells in the PPE_central_. This crosstalk consists of Dpp and additional unidentified signals that control peripodial BM_PPE_ growth and mechanics, potentially by controlling matrix remodeling via regulation of MMP2 expression (see below).

## Discussion

In this work we uncover the emergence of cell morphological asymmetry within a growing epithelial organ and characterize the underlying mechanics. This asymmetry arises from the spatial regulation of cell growth and proliferation, differential mechanical resistance of the BM and tissue scale force transmission. Distinct cell shape transitions are the result of elastic deformations determined by tissue-internal and -external stresses and local material properties. In the wing disc, cuboidal-to-columnar transitions arises from growth-induced (tissue-intrinsic) compressive stress dependent on elastic BM properties that resist planar growth. In contrast, cuboidal-to-squamous shape changes arise from cell stretching by a tissue-extrinsic tensile stress dependent on stressless BM growth that allows stress transmission onto the cell layer.

We initially focused on the origin of the forces inducing epithelial cells shape transitions in the PPE. We identified an asymmetry in growth between the DP and the PPE layers and a planar asymmetry in proliferation within the PPE layer. While previous work proposed that growth in the DP and the PPE are coordinated^24^, we find a growth asymmetry between the two layers (higher growth in the DP, see Figure S2), which leads to tensile stress in the slower growing PPE layer. This tensile stress is further amplified by DP bending leading to additional area expansion and elastic flattening of squamous cells (Figure 4). Therefore, tissue-extrinsic, tensile stress arising from a growth asymmetry between tissue layers and DP bending induce the cuboidal-to-squamous cell shape transition during peripodial morphogenesis. Notably, mechanical modelling suggested that squamous flattening of the anterior follicle epithelium during *Drosophila* oogenesis^51^ is a response to tensile stress originating from the faster growing underlying germline. In addition, engineering approaches hint that the differentiation of the squamous epithelium in the alveoli of our lungs depends on tensile forces^52^ and stretching forces were associated with the squamous morphogenesis of the retinal pigment epithelium during eye development^16^. Therefore, tissue-extrinsic tensile forces may be a conserved mechanism driving cell flattening during squamous morphogenesis.

We found that this tensile force mechanically leads to spatially distinct tissue and cell deformations: cuboidal-to-columnar in the hinge and cuboidal-to-squamous in the central cells. We first investigated the stresses present in the peripodial cell layer. As DP bending leads to stretching of the PPE layer, we expected the whole peripodial cell layer to be under spatially uniform tensile stress. However, we found that only the central squamous epithelium is under *tensile* elastic stress, indicated by in-plane relaxation of the tissue and a concomitant increase in tissue thickness upon the loss of disc bending (see Fig.4). In contrast, we observed *compressive* elastic stress in the peripodial hinge cells (hinge_A_ and Hinge_P_), indicated by tissue flattening upon acute removal of the BM_PPE_. Importantly, we find no difference in the growth rate of peripodial hinge and squamous cells (Figure 3E) indicating that spatial differences in growth rate cannot account for differential cell deformations in the PPE.

However, we revealed that differential regulation of the BM_PPE_ is key for the distinct stress pattern in the epithelial layer and the resulting cell shape transitions. While the BM_PPE_ is stressless, i.e. relaxed in central squamous cells, it is stretched and under tensile stress in the hinge domain (Figure 5). Notably, the thick BM_PPE_ of the hinge shields cells from the tissue-extrinsic tensile stress, leading to the accumulation of compressive elastic stress in the cell layer and its cuboidal-to-squamous transition. In contrast, squamous morphogenesis requires a thin BM_PPE_ that relaxes external tensile stress, i.e. that remains stress-free, thereby transducing tensile stresses onto the epithelial layer which leads to cell flattening.

These spatial differences in the BM_PPE_ could be due to differences in BM stiffness or a result of differential regulation of BM growth (or a combination of both). Indeed, previous work has shown that mutations in the BM component Perlecan (Trol in *Drosophila*) lead to increased BM stiffness and inhibit the flattening of the PPE_central_ cells^29^ (note that discs also fail to bend in this scenario). These results suggest that Trol is required to soften the BM, so they cannot fully explain the observed differences in the BM_PPE_ stress pattern: First, Trol levels are higher in the BM_PPE_ of the hinge than in the squamous cells, which should increase stiffness in the central BM_PPE_, and thereby resist planar expansion of overlying cells, contrary to observations. Second, we find that the BM_PPE_ covering the squamous cells is effectively stress-free (see Figure 5 and ^19,29^), i.e. not elastically deformed. This last observation highlights that the BM_PPE_ of squamous cells is not just potentially softer but that it grows at the same rate as the tissue expands, in other words its reference state increases as it grows. Hence, the more likely mechanism is a differential regulation of BM growth properties. We have previously shown that the BM_PPE_ of squamous cells follows a planar mode of growth^19^. Importantly, planar BM growth will lead to an increase in the rest state of the BM_PPE_ as the squamous cells are stretched, such that its rest configuration adjusts to tissue size (see Figure S10). This scenario does not require differences in the material properties of the BM (such as stiffness) but depends on local regulation of BM growth orientation. While the regulation of BM growth remains not well understood, active matrix remodeling via MMP2 represents a potential mechanism of controlling BM growth orientation. This is supported by our findings that squamous cells express high levels of MMP2 and any reduction in MMP2 levels impairs squamous morphogenesis (Figure S9A-C and ^19,29^).

In addition, we uncover an asymmetry in the mechanical coupling and force transmission between the epithelial layers of the wing disc. We identify Dpy^39^ as a direct mechanical connection between the apical surface of the two epithelial layers of the wing disc. Notably, global disc geometry and bending result in the proximity of DP and PPE only in its central region (Figure S6C), leading to the selective shearing of squamous cells. Dpy was shown to act as a physical connector between the pupal cuticle and epithelial^41^ and tendon cells^42^ and between the chitinous extracellular matrix and the tube cells of the tracheal network^53^. Therefore, our findings reveal also epithelial-epithelial coupling to Dpy functions during *Drosophila* development.

Finally, we find that Dpy is required for contact-induced signaling that alters BM_PPE_ mechanics and epithelial morphology via creating an asymmetry in MMP2 expression (low in the DP and high in the PPE, Figure S9A). Loss of Dpy from the surface of both, the DP and PPE tissue layer results in decreased peripodial MMP2 expression (Figure S9D) and in addition leads to ectopic Dpp signaling. These results suggest that Dpy possibly controls paracrine signaling between the two tissue layers. Notably, the large extracellular domain of Dpy contains over 300 EGF-repeats^39^ which are known to be glycosylated and to mediate protein-protein interactions^54^. Hence, Dpy could bind paracrine factors such as Dpp in the luminal space of the wing disc and act as ‘insulator’ between the two epithelial layers. However, ectopic activation of Dpp signaling is not sufficient to promote the observed increase in cell height (see Figure S8E-F). Therefore, additional signaling factors might be involved. Wingless, another paracrine factor that is also expressed at high levels in the DP^55^ was shown to induce cuboidal cell shape when overexpressed in the squamous cells of the wing disc^25^. An alternative, attractive hypothesis is mechano-signaling between the PPE and DP epithelial layers as Dpy-mediated coupling leads to stress in the PPE. MMP2 expression could be controlled by stress-induced signaling via nuclear compression of PPE cells^56,57^, mechano-sensitive channels^58^ and/or via the Hippo/YAP pathway^59^. Overall, our results highlight Dpy as part of a signaling network that orchestrates multilayer morphogenesis and bending by setting up an asymmetry in mechanical properties between the tissue layers.

Together, this work shows that epithelial tissue morphogenesis emerges from cellular and tissue level interactions coupled by mechanical and geometric feedback through the basement membrane and Dpy. BM properties through Collagen IV and Dpy as an inter-tissue mechanical linker play a pivotal role in the regulation of both force generation and transmission across cell and tissue scales as the organ grows. It will be exciting to see how future research will shed light on the intricate regulation of matrix mechanics and growth in the making of an organ.

## Acknowledgments

We would like to thank all the members of the Lecuit team for stimulating discussions. We thank Benoît Aigouy (IBDM, Marseille, France), Markus Affolter (Biozentrum, Basel, Switzerland), Kendal Broadie (Vanderbilt, USA), Loic LeGoff (Institut Fresnel, Marseille, France), Stéphane Noselli (IBV, Nice, France), Frank Schnorrer (IBDM, Marseille, France), the Bloomington *Drosophila* Stock Center and the Developmental Studies Hybridoma Bank (University of Iowa, USA) for flies and reagents. The IBDM imaging platform and the France-BioImaging infrastructure supported by the Agence Nationale de la Recherche (ANR-10-INSB-04-01; call “Investissements d’Avenir”) provided support. This work was supported by the Ligue Nationale Contre le Cancer (Equipe Labellisée 2018). S.H. was supported by an ATER position from the College de France (Paris, France) and the Wellcome Trust (Career Development Award 227832/Z/23/Z).

## Author contributions

S.H. and T.L. conceived and designed the study. S.H. performed the experiments and quantified the data. S.H. wrote the first draft of the manuscript, T.L. made comments and edited the text. All authors read, commented, and approved the manuscript.

## STAR Methods

### RESOURCE AVAILABILITY

#### Lead contact

Further information and requests for resources and reagents should be directed to and will be fulfilled by the lead contact, Thomas Lecuit. A source data file is provided with the paper.

#### Materials availability

This study did not generate new unique reagents.

#### Data and code availability

- Microscopy data reported in this paper will be shared by the lead contact upon request.
- The Python code to visualize epithelial height will be shared by the lead contact upon request.
- Any additional information required to reanalyze the data reported in this paper is available from the lead contact upon request.

### EXPERIMENTAL MODEL DETAILS

#### Drosophila strains used

The following fly lines were used: *y*^1^*,w*^1118^, *hs-Flp; act > Stop > Gal4, UAS-EGFP* (*AyGAL4*, originating from Bloomington stock #64231), *hh-Gal4* ^60^, *Vkg^G^*^454^*::GFP* ^61^ (from F. Schnorrer), *UAS-MMP2* (obtained from L. Le Goff, originally from the M. Grammont lab), *Dpy::YFP* ^47,48^ (from Benoît Aigouy), UAS-Dad ^62^ (on 2nd) and UAS-Tkv1A (on 2nd, both from M. Affolter). The following lines were obtained from the Bloomington stock center: *AGiR-Gal4* (#6773), *UAS-CD8::mRFP* (#27398), *UAS-CD8::RFP* (#27392), *sqh-Sqh::GFP* (#57144), *tsh-Gal4* (#3040), *tub-Gal80^ts^* (#7017), UAS-Myc.Z (#9674), UAS-Mnt / CyO (#80569), UAS-DpyTRiP (#36691).

#### Genotypes by Figure

- Figure 1: *w*^1118^
- Figure 2: *hs-Flp; act > Stop > Gal4, UAS-EGFP*
- Figure 3: (**A-B, F**) w118, (**C**) *hs-Flp; act > Stop > Gal4, UAS-EGFP*
- Figure 4: *Vkg::GFP / sqh-Sqh::GFP*
- Figure 5: *Vkg::GFP / tsh-Gal4 ; UAS-CD8::mRFP / +*
- Figure 6: (**B**) *Dpy::YFP* (**D-F**) control: *UAS-CD8::RFP / + ; AGiR-Gal4 / +* | PPE_KD_: *UAS-CD8::RFP / + ; AGiR-Gal4 / UAS-DpyTRiP*
- Figure 7: (**A**) control: *UAS-CD8::RFP / + ; hh-Gal4 / +* | Posterior_KD_: *UAS-CD8::RFP / + ; hh-Gal4 / UAS-DpyTRiP* (**C**-**F**) *UAS-CD8::RFP / Vkg::GFP ; hh-Gal4 / UAS-DpyTRiP*
- Supplementary Figure 1: (**A**) *w*^1118^ (**D**) *hs-Flp; act > Stop > Gal4, UAS-EGFP*
- Supplementary Figure 3: (**A-C**) *Vkg::GFP / sqh-Sqh::GFP* (**D-F**) Vkg::GFP / Ecad::GFP
- Supplementary Figure 4: (**A-G**) control: *nub-Gal4 / + ; tub-Gal80ts / UAS-CD8::mRFP* | MMP2_OE_: *nub-Gal4 / UAS-MMP2 ; tub-Gal80ts / UAS-CD8::mRFP* (**H**) *Vkg::GFP / tsh-Gal4 ; UAS-CD8::mRFP / +*
- Supplementary Figure 5: (**A**) Dpy::YFP (**C**) control: Dpy::YFP / + | PPE_KD_: *UAS-CD8::RFP / Dpy::YFP ; AGiR-Gal4 / UAS-DpyTRiP* | Posterior_KD_: *UAS-CD8::RFP / Dpy::YFP ; hh-Gal4 / UAS-DpyTRiP*
- Supplementary Figure 6: (**A-D**) control: Dpy::YFP / + | PPE_KD_: *UAS-CD8::RFP / Dpy::YFP ; AGiR-Gal4 / UAS-DpyTRiP* | Posterior_KD_: *UAS-CD8::RFP / Dpy::YFP ; hh-Gal4 / UAS-DpyTRiP* (**E**) | control: *UAS-CD8::RFP / Dpy::YFP ; UAS-DpyTRiP / +* | PPE_KD_: *UAS-CD8::RFP / Dpy::YFP ; AGiR-Gal4 / UAS-DpyTRiP*
- Supplementary Figure 7: control: *UAS-CD8::RFP / + ; hh-Gal4 / +* | Posterior_KD_: + */ Dpy::YFP ; hh-Gal4 / UAS-DpyTRiP*
- Supplementary Figure 8: (**A-B**) control: *UAS-CD8::RFP / + ; AGiR-Gal4 / +* | PPE_KD_: *UAS-CD8::RFP / + ; AGiR-Gal4 / UAS-DpyTRiP* (**C-D**) control: *UAS-CD8::RFP / nub-Gal4* | DP_KD_: *UAS-CD8::RFP / nub-Gal4 ; UAS-DpyTRiP / +* (**E-F**) control: *UAS-CD8::RFP / + ; AGiR-Gal4 / +* | Dad_AGiR_: *UAS-CD8::RFP / UAS-Dad ; AGiR-Gal4 / +* | TkvCA_AGiR_: *UAS-CD8::RFP / UAS-Tkv1A ; AGiR-Gal4 / +*
- Supplementary Figure 9: (**A-D)** control: UAS-CD8::RFP / + ; hh-Gal4 / + | Posterior_KD_: *UAS-CD8::RFP / + ; hh-Gal4 / UAS-DpyTRiP* (**E**) *UAS-CD8::RFP / Vkg::GFP ; hh-Gal4 / UAS-DpyTRiP*

#### Staging and sample collection

All data shown originates from staged tissues with developmental time indicated in hours after egg laying (hAEL) as described in Harmansa et.al.^19^. In brief, flies were allowed to deposit eggs for a two-hour interval in standard fly tubes supplemented with yeast paste. Collected embryos were subsequently kept in an incubator at 25°C and larvae dissected at the desired developmental stage. At 25°C the second to third instar transition corresponds to 72hAEL and the end of 3^rd^ instar development to 120hAEL. For the timepoints older than 72hAEL only male larvae were included in the study, given the differences in growth between male and female larvae. Male larvae were positively selected by the presence of two translucent spots in the posterior half of the larvae, corresponding to the male genitalia disc. These spots are not clearly visible at 72hAEL and before and hence these timepoints include male and female samples.

All samples of one experiment were dissected together using the same solutions (fixative, PBS, antibody dilutions etc.) to reduce experimental variations and to allow direct comparison between different developmental stages.

### METHOD DETAILS

#### Antibodies

Primary antibodies listed in the Key Resource Table were used at the following concentrations: Mouse-anti-Wingless (4D4-s) at 1:120; Mouse-anti-Patched (Apa1-s) at 1:40; Rat-anti-DE-cadherin (DCAD2 concentrate) at 1:200; Rabbit-anti-GFP (ab6556) at 1:1000; Rabbit-anti-Phospho-Histone H3 (PHH3) at 1:1000; Rabbit-anti-MMP2 at 1:500.

Tissue outlines were marked by Alexa Fluor 660 Phalloidin, used at 1:100 and added with the secondary antibodies. For labeling DNA content DAPI dye was added during the secondary antibody incubation period (stock solution at 5mg/ml, used at 1/1000 dilution). Secondary antibodies used at 1/500 dilution were Alexa 488 (Donkey anti-mouse A21202, Donkey anti-rabbit A21206), Alexa 568 (Donkey anti-mouse A10037, Donkey anti-rabbit A10042, Goat anti-rat A11077) and Alexa 647 (Donkey anti-rabbit A31573) from Invitrogen and Alexa 647 (Donkey anti-mouse 715 605 151) from Jackson ImmunoResearch. We used 2% normal donkey serum as a blocking agent (017-000-121, Jackson ImmunoResearch).

#### Immunohistochemistry and imaging

Imaginal disc tissues were stained as described previously^63^. Samples were mounted in Vectashield Plus (H-1900, Vector Laboratories). To conserve natural tissue morphology and avoid squishing, samples were mounted using double-sided tape as spacers (TESA 05338). All fixed samples were imaged on a Leica SP8 confocal microscope (40x or 63x/1.4NA oil immersion objectives). All data for one experiment was acquired within the same imaging session under identical microscope settings, ensuring comparability of fluorescence intensities. For optical cross sections, stacks with high z-axis resolution were acquired (0.33μm slice spacing).

#### Induction of cytosolic GFP clones

Related to Figure 2, Figure 3C and Figure S1D. Stop-cassette removal was induced by heat shock at 37°C leading to the expression of cytosolic GFP and Histone::RFP (hs-Flp; act > Stop > Gal4, UAS-EGFP/UAS-Histone::RFP). In general, heat-shock was induced 24 hours before dissection and heat shock length was adjusted to the developmental stage to obtain sparse clone induction (3 min at 48hAEL, 4 min at 72hAEL, 7 min at 96hAEL).

#### ex vivo culture

Related to Figure 3C-E. The used *ex vivo* culture conditions are based on the protocol published by Dye *et.al.* ^64^ with the following modifications: Culture medium was supplemented with adenosine deaminase (ADA, 8.3ng/ml final concentration, Roche 10102105001) as described previously ^19^.

Wing discs from *hs-Flp; act > Stop > Gal4, UAS-EGFP/UAS-Histone::RFP* larvae, were dissected in culture medium 24 hours after heat shock (as described in the previous section). Explants were immobilized under a permeable membrane Whatman cyclopore polycarbonate membranes; Sigma, WHA70602513) using double sided tape as spacer (∼90 μm thickness, Tesa 05338, Reichelt). The mounted samples were inserted in an Attofluor chamber (A7816, ThermoFisher), immersed in 1 ml of culture medium and covered by parafilm to reduce evaporation during long-term imaging. Explant movies were acquired on a Nikon Roper spinning disc Eclipse Ti inverted microscope (controlled by Metamorph 7.8.4.0) using a 40x/1.25 N.A. water immersion objective at 22 °C. Stacks of 1 μm *z*-spacing were acquired every 10 min for up to 12 h.

#### Collagen IV digestion and MyoII inhibition (acute and genetic)

Related to Figure 4 and Figures S3-4. For acute Collagen IV digestion and MyoII inhibition, larvae of genotype *Vkg::GFP*^454^ */ sqh-Sqh::GFP* were dissected in PBS at indicated time points. Dissected larvae were transferred to 2 ml Eppendorf tubes containing PBS at 37 °C on a heat block. The ROCK inhibitor Y-27632 dihydrochloride (Sigma-Aldrich, Y0503) was added to 2 mM final concentration for 2 min to inhibit MyoII activity. Subsequently, Collagenase (Sigma-Aldrich, C0130) was added to a final concentration of 1 mg/ml and incubated for 1 min to digest the basement membrane. After one minute fixative (4% PFA in PBS) was added to conserve tissue morphology. Discs were fixed for 20 min at RT on a shaker and then processed for immunostaining as described above.

For genetic digestion of the disc proper basement membrane MMP2 (UAS-MMP2) was over-expression specifically in the disc proper epithelium using *nubbin-Gal4* (*nub-Gal4*). Temporal expression of MMP2 was controlled using a temperature sensitive version of Gal80 (Gal80^ts^). To avoid leaky expression of MMP2 (and the resulting lethality) experiments involving *nub-Gal4 / UAS-MMP2 ; tub-Gal80ts / UAS-CD8::mRFP* larvae were performed at RT. At mid-3rd instar stage the larvae were shifted to restrictive temperature (29 °C) to induce overexpression of MMP2. After 7 hours at 29 °C larvae were dissected in PBS and exposed to Y27632 to inhibit MyoII activity as described above. Subsequently, the tissues were fixed and processed for immunostaining to detect Vkg levels (Rabbit-anti-Vkg, 1:500, from Van de Bor ^65^).

#### Circular bleaching and decellularization essay

Related to Figure 5, Figure 7D-F, Figure S4H and Figure S9E. The circular bleaching and decellularization essay for wing disc explants is described in detail in Harmansa *et.al*. ^19^. In brief, wing discs of staged larvae (expressing *Vkg::GFP*^454^) were isolated in PBS and immobilized by gluing (using classical embryo glue) their notum to the bottom of a petri dish in a drop of PBS, with the PPE facing upwards. Mounted discs were imaged on an upright Nikon A1R MP+ multiphoton microscope (controlled by Nikon NIS-Elements AR 5.11.01 software) using a water-immersion objective (40x/1.15NA). A tunable wavelength pulsed laser (Coherent) at 920 nm was used to excite Vkg::GFP. Circular regions of interest (ROI) were bleached on the Vkg::GFP signal of the PPE layer in the central region (PPE_center_) and in the anterior and posterior peripodial hinge region (Hinge_A_ and Hinge_P_, respectively). Bleaching of ROIs was limited to a maximum of 40 min, allowing to image ∼4-5 discs. After imaging the marked ROIs, the discs were exposed to 3% Triton X-100 in PBS to decellularize and remove the tissue layers. After a 60 min incubation at RT, the ROIs were re-imaged in their relaxed configuration. Changes in the ROI area Vkg::GFP intensity were quantified as described in the Quantification section.

### QUANTIFICATION AND STATISTICAL ANALYSIS

#### Image processing

Raw data image processing was performed in Fiji/ImageJ software and subsequent quantifications were performed in Python.

##### Quantification of apical cell surface area along the A/P-axis

We obtained high resolution image stacks of staged wing discs stained for E-cadherin (Ecad), outlining the apical cell surface, and Wingless/Patched (Wg/Ptc), providing landmarks. Given a lack of suitable landmarks in the peripodial layer, we used the landmarks provided by the Wg/Ptc staining in the DP layer: In this study we focus on peripodial morphology in a rectangular region spanning the full anterior-posterior (A/P) axis of the disc and slightly dorsal to the DP D/V-boundary (marked by the Wg signal).

We used the Fiji plugin ‘LocalZProjector’ ^66^ to obtain 2D surface projections and height maps of the peripodial apical surface (Ecad signal). Using these projections, apical cell area was segmented using the Fiji ‘Tissue Analyzer’ plugin ^67^. Subsequently, we corrected for the introduced error in surface area due to the 2D projection by using the DeProject Matlab tool ^66^: DeProjects utilizes the 2D segmentation output from Tissue analyzer and the height map from LocalZProjector to effectively deproject the 2D segmentation to 3D, thereby correcting for curvature or tissue bending induced errors. The corrected apical surface area was plotted along the A/P-axis (as shown in Figure 1F), with *x*=0 corresponding to the DP A/P-boundary (marked by the Ptc staining). Line plots were created using the seaborn.lineplot function of the Seaborn package ^68^ in Python, error bands depict the standard deviation.

##### Epithelial height quantification along the A/P-axis

Epithelial height was quantified from optical cross sections of wing disc stained for Wg/Ptc (+ Phalloidin in few cases) parallel to and just dorsal to the DP D/V-boundary (using the ‘Reslice [/]’ function in Fiji). In these cross-sections, the PPE apical and basal surface (sufficiently labeled by Wg/Ptc) were marked by spline-fits using the Fiji plugin ‘Kappa - Curvature analysis’ ^69^ (version 2.0.0). The apical and basal surface spline-fits were exported and further processed in Python. Using a custom script, we computed the local minimal distance between the two spline-fits, corresponding to the local height of the PPE. For height plots (as shown in Figure 1G), PPE height was subsequently plotted along the A/P-axis, *x*=0 corresponding to the DP A/P-boundary (marked by Ptc), using the seaborn.lineplot function of the Seaborn package ^68^ in Python. In addition, we used a custom Python script to visualize local epithelial height by a color-code (blue to red for minimal to maximal height) onto the cross-section images (as shown in Figure 1E). In these images, the apical and basal surface spline-fit are indicated by dashed lines and the local PPE height by solid-colored bars.

##### Quantification of cellular aspect ratio

To visualize developmental changes in cell shape, we have adapted a framework to quantify the cellular aspect ratio during peripodial morphogenesis (see for example Figure 1H). While in cuboidal epithelia the ratio of apical cell diameter (*D*) to cell height (*H*) is roughly on, this ratio is >1 in squamous and <1 in columnar epithelia (see Figure 1A). While epithelial *H* was directly measured from cross-section images (see above), we approximated the apical cell surface as a circle to obtain D 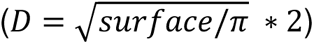. For plotting the average cell shape ratio (*D/H*) along the A/P-axis we calculated *D* for individual cells and divided it by the average local height *H* for a given genotype and time point.

##### Cell volume quantification

Related to Figure 2. We acquired image stacks with high z-resolution (slice distance = 0.33µm) of wing discs expressing cytosolic GFP and Histone::RFP in clones (hs-Flp; act>Stop>Gal4, UAS-EGFP/UAS-Histone::RFP) stained for anti-GFP (clone volume) and anti-Wg/Ptc (landmark). Individual GFP-positive clones were cropped in Fiji/ImageJ, background subtracted (‘rolling ball’ function with radius of 50px), and the clone volume segmented in Ilastik ^70^ using the ‘pixel classification’ mode. Clone volume was divided by cell number per clone, obtained by the Histone::RFP signal and plotted against the position along the A/P-axis (*x*=0 corresponding to the DP A/P-boundary).

##### Quantification of mitotic rate using PHH3 staining

Related to Figure 3A-B. Wing discs of *y*^1^*, w*^1118^ larvae were dissected at indicated time points and stained for E-cadherin (cell outlines), Phospho-HistoneH3 (PHH3, mitotic cells) and Wingless/Patched (Wg/Ptc, landmarks to register multiple discs). Projections of the peripodial apical surface (marked by Ecad) were obtained using the Fiji plugin ‘LocalZProjector’ ^66^ as described above. Projections were segmented using the Fiji ‘Tissue Analyzer’ plugin ^67^ and mitotic cells marked by hand using the ‘Multi-point’ selection tool in Fiji. The subsequent analysis was performed in Python: Using the landmarks provided by the Wg/Ptc cross and the Wg ring, the cell and mitotic data was registered between individual discs using affine transformation. We then computed the average mitotic rate for three distinct regions (see Figure 3B), the central PPE (PPE1), and the circular intermediate region (PPE2) and the peripheral hinge region (Hinge). Subsequently, we obtained the mitotic rate by dividing the number of mitotic cells (PHH3) by the average cell number in the respective region. This calculation was done for all three regions of each disc (see individual points in Figure 3B, *right*) at the indicated time points.

##### Growth rate quantification in ex vivo cultured discs

Related to Figure 3C-E. We performed volumetric *ex vivo* imaging of 96hAEL old wing disc, expressing cytosolic GFP in a clonal manner for up to 12 hours (10 min imaging intervals, as described in Harmansa et.al.^19^). Individual GFP-positive clones were cropped in Fiji/ImageJ, background subtracted (‘rolling ball’ function with radius of 50px), and the clone volume segmented in Ilastik ^70^ using the ‘pixel classification’ mode. Clone volume was quantified for every hour of a movie by averaging the three consecutive volumes and normalized by the initial volume to plot the relative volume over culture time (see Figure 3D). In addition, the hourly growth rate was calculated by:

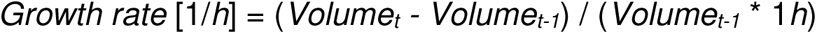

where *Volume_t_* indicates the clone volume at *time* = *t*, and *Volume_t-1_* the clone volume one hour earlier. We subsequently calculated an average growth rate for each clone (blue dots in Figure 3E) and an overall growth rate for peripodial cells after 96hAEL.

##### Nuclear DAPI intensity quantification

Wing discs of *y*^1^*, w*^1118^ larvae at indicated time points were stained for their DNA content using a DAPI staining. The fluorescence intensity of the DAPI staining is largely proportional to DNA levels and hence, the integrated DAPI fluorescence per nucleus is a good measure for the cellular DNA content. We have segmented individual nuclear volumes using the 3D-Cell-Annotator plugin ^71^ for the Medical Imaging Interaction Toolkit (MITK) to obtain nuclear masks. Using these masks on the DAPI channels allowed us to extract the integrated nuclear DAPI intensities of the PPE_central_ and the Hinge region at defined time points (see Figure 3G). The integrated DAPI intensity was plotted in respect to clone position (either in the center or periphery, PPE_central_ or Hinge,respectively).

##### Collagen IV intensity quantifications

Related to Figure S4F: Intensity of the Collagen IV along the basal surface of the PPE was quantified from cross-section images of discs stained for Vkg and Wg/Ptc. The basal surface visualized by the Wg/Ptc signal was marked by a spline-fit (using the Fiji/ImageJ plugin ‘Kappa - Curvature analysis’ ^69^ (version 2.0.0) and Vkg fluorescence was extracted along this line and plotted in respect to the A/P-axis (*x*=0 corresponding to the DP A/P-boundary).

Related to Figure 7C: Here Vkg::GFP fluorescence was quantified in Fiji/ImageJ using the ‘Segmented line’ tool marking the basal outline and a line width that covered the Vkg::GFP signal.

##### Quantification of basal disc outline

Related to Figure S4D and Figure S6B: The average disc proper (DP) basal outline of a genotype was computed using the ‘Kappa - Curvature Analysis’ ^69^ plugin in Fiji as described in Harmansa *et.al*. ^19^. In brief, the basal outline of the DP was marked by a spline-fit, and subsequently multiple discs were aligned according to the position of the DP A/P boundary (outlines of the individual discs are shown as thin lines in Figure S4D). An average spline fit (indicated by thick) was computed in Python.

##### Quantification of distance between DP and PPE apical surface

Related to Figure S6A+C: To quantify the distance between the apical surface of the PPE and the DP epithelium we re-used the approach described in the ‘Epithelial height quantification along the A/P-axis’ section. Instead of marking the apical and basal surface of the PPE by a spline fit, we marked the apical surface of the PPE and the DP with spline-fits and computed and visualized the distance between the two epithelial layers as described before.

##### Assessing stress in the BM using decellularization essays

Related to Figure 5C and Figure 7D-E: As previously described in Harmansa et.al. ^19^, we acquired image stacks of the bleached, circular regions of interest (ROI) before and 60 min after decellularization. The obtained image data was oriented and projected such that the circular ROIs were in-plane and not tilted (which would lead to an error in area quantification, using the ‘reslice’ and ‘transformation’ functions in Fiji/ImageJ). Using these projections, the area of each ROI was measured before and after decellularization in Fiji using the ‘polygon selection’ tool. Relative changes in ROI area upon decellularization were plotted in Python (Seaborn library).

##### Quantifying BM thickness in decellularization essays

Related to Figure 5D and Figure 7F: As previously described in Harmansa et.al. ^19^, we extracted the Vkg::GFP intensity in cross-section views (average projections of 5 slices) using the ‘straight line tool’ (10px wide, corresponding to 6.2μm) and the plot profile function. For each disc, Vkg::GFP intensity was quantified at three positions (close to the bleached ROIs) and averaged. Profiles were aligned by their peak intensity to obtain average profiles, plotted in Python using the Seaborn ‘lineplot’ command (as shown in Figure 5D). To quantify BM thickness, we used intensity thresholds to determine the width of the Vkg::GFP intensity profile. Thresholds were chosen for each experiment to capture the majority of the Vkg::GFP profile (threshold value 120 in Figure 5D and 40 in Figure 7F). Notably, thresholds vary between experiments due to differences in imaging conditions.

##### P-Mad profiles in the DP and the PPE

Related to Figure 6B and Figure 7B+D: The spatial signaling activity of Phospho-Mad (P-Mad) was computed from cross-section images of discs stained for P-Mad and Wg/Ptc. Profiles were extracted using the ‘Segmented line’ tool in Fiji/ImageJ either in the PPE or the DP layer, as indicated in Figure S6B, *top*. Individual profiles were registered to the DP A/P-boundary (defined as *x*=0) and averaged per genotype. Profiles were plotted in Python using the Seaborn package (‘lineplot’ command), the error band indicating the standard deviation.

##### Quantification of mean MMP2 levels

Related to Figure S9: We obtained cross-section images of wing discs stained for MMP2 and Wg/Ptc. We then quantified MMP2 levels along the A/P axis in Fiji/ImageJ, using the ‘segmented line’ tool (10px line width, corresponding to 3.2 μm) for Figure S9B-C or using the ‘Polygon selection’ tool to mark the anterior and the posterior tissue regions of the PPE and quantify the mean MMP2 fluorescence intensity (as indicated in Figure S9D, *top*).

Intensity profiles along the A/P-axis were aligned (A/P_DP_ corresponding to x=0) and plotted together with the epithelial height in Phyton (Seaborn package, ‘lineplot’ command, error band indicating the standard deviation). We subsequently quantified the average MMP2 intensity and epithelial height for each disc in the Hinge_A_ (purple), PPE_central_ (blue) and Hinge_P_ (green, see indications in Figure S9B) and plotted average height against MMP2 levels (Figure S9C, Python, Seaborn package using the ‘regplot’ command). Statistics of the linear regression were obtained using the Python scipy.stats package (‘pearsonr’ and ‘linregress’ command).

Average MMP2 intensity in the anterior and posterior hinge region (as indicated in Figure S9D, *top*) was plotted as boxplots in Python (Seaborn package, ‘boxplot’ command).

### Statistics

#### Data representation

Data was plotted in Python (3.9.7) using the Seaborn library (0.11.2). In line plots, the error bands indicate the standard deviation. In box plots, the median is indicated by a central thick line while the interquartile range (containing 50% of the data points) is outlined by a box. Whiskers indicate the minimum and maximum data range; outliers are indicated by a black rhomb and were excluded from further processing.

#### Statistical tests and significance

Given the experimental constraints we aimed to obtain a sample size large enough (*n* ≥ 5) to allow testing statistical significance by using a two-sided Student’s *t* test (unequal variance, *P ≤ 0.05, **P ≤ 0.005, ***P ≤ 0.0005). The number of samples and P-values are either indicated in the figure or the respective legend. For each experiment, *n-*numbers indicate biological replicates, meaning the number of biological specimens evaluated (e.g., the number of wing discs or clones). Statistical significance was assessed using a two-sided Student’s *t*-test (unequal variance, in MS Excel, *P ≤ 0.05, **P ≤ 0.005, ***P ≤ 0.0005) when comparing two groups. For comparing more than two groups, statistical significance was assessed using a one-way ANOVA and Tukey’s post hoc test (\**p* ≤ 0.05, \*\**p* ≤ 0.005, \*\*\**p* ≤ 0.0005, in Python, statsmodels.stats.multicomp package using the ‘pairwise_tukeyhsd’ command). Linear regression and correlation analysis was performed in Python using the scipy.stats package (‘pearsonr’ and ‘linregress’ command).

### KEY RESOURCES TABLE

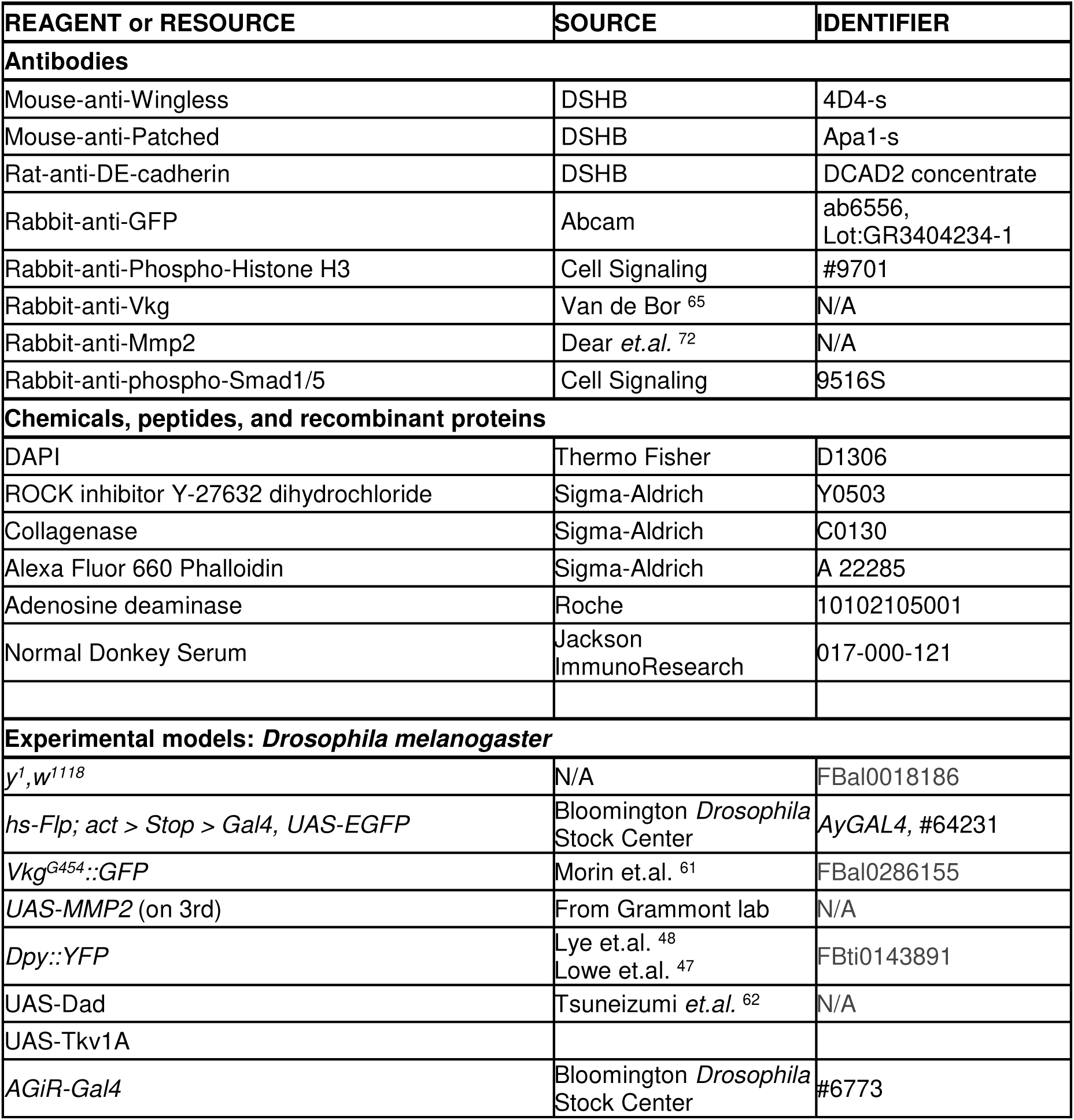
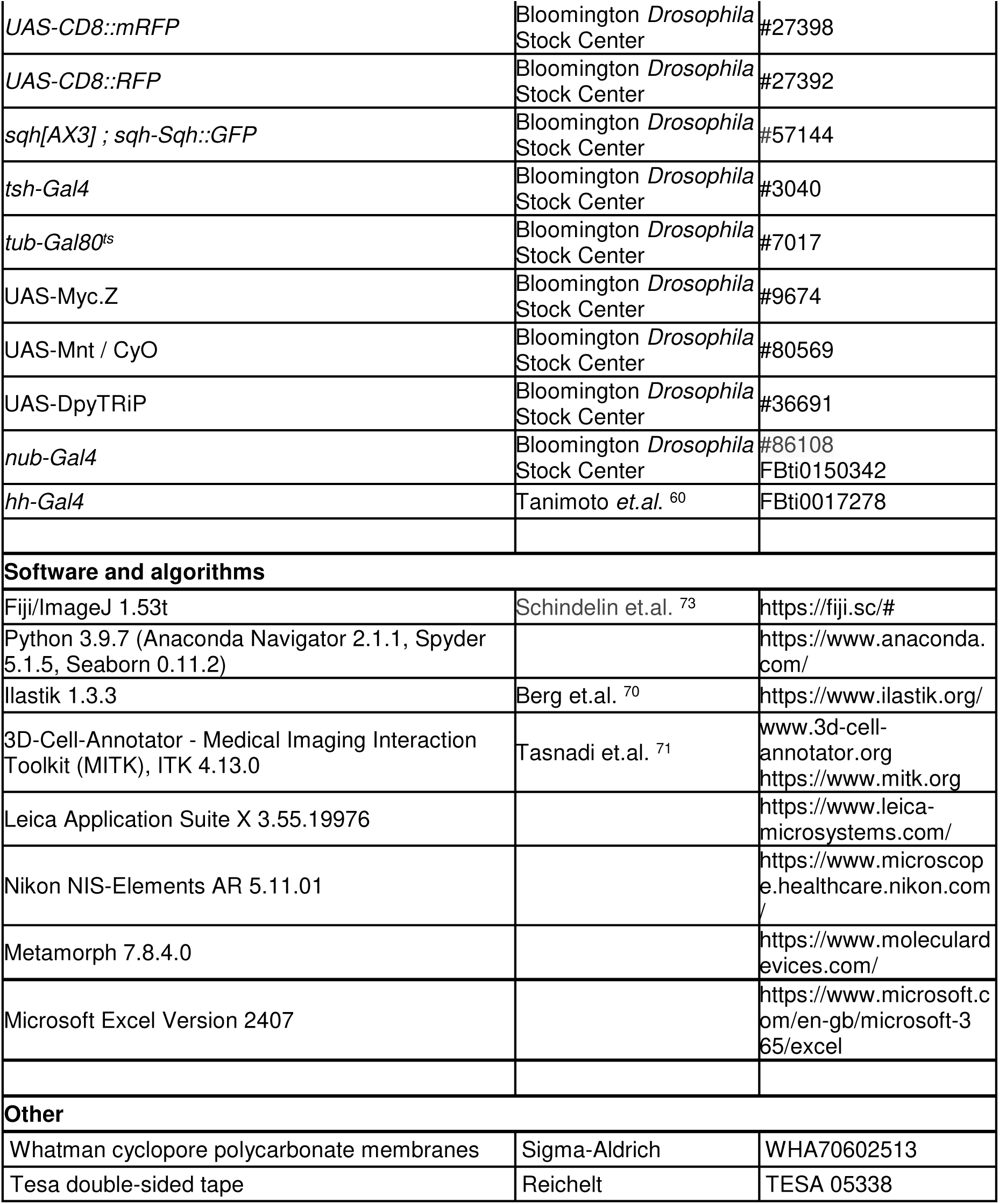

## Bibliography

1. Collinet, C., and Lecuit, T. (2021). Programmed and self-organized flow of information during morphogenesis. Nat. Rev. Mol. Cell Biol. 22, 245–265. 10.1038/s41580-020-00318-6.

2. Agarwal, P., and Zaidel-Bar, R. (2019). Principles of Actomyosin Regulation In Vivo. Trends Cell Biol. 29, 150–163. 10.1016/j.tcb.2018.09.006.

3. Stooke-Vaughan, G.A., and Campàs, O. (2018). Physical control of tissue morphogenesis across scales. Curr. Opin. Genet. Dev. 51, 111–119. 10.1016/j.gde.2018.09.002.

4. Savin, T., Kurpios, N.A., Shyer, A.E., Florescu, P., Liang, H., Mahadevan, L., and Tabin, C.J. (2011). On the growth and form of the gut. Nature 476, 57–62. 10.1038/nature10277.

5. Thompson, D.W. (1992). On growth and form (Dover).

6. Vignes, H., Vagena-Pantoula, C., and Vermot, J. (2022). Mechanical control of tissue shape: Cell-extrinsic and -intrinsic mechanisms join forces to regulate morphogenesis. Semin. Cell Dev. Biol., S1084952122000842. 10.1016/j.semcdb.2022.03.017.

7. Schöck, F., and Perrimon, N. (2002). Molecular Mechanisms of Epithelial Morphogenesis. Annu. Rev. Cell Dev. Biol. 18, 463–493. 10.1146/annurev.cellbio.18.022602.131838.

8. Díaz-de-la-Loza, M.-C., and Stramer, B.M. (2024). The extracellular matrix in tissue morphogenesis: No longer a backseat driver. Cells Dev. 177, 203883. 10.1016/j.cdev.2023.203883.

9. Wu, D., Yamada, K.M., and Wang, S. (2023). Tissue Morphogenesis Through Dynamic Cell and Matrix Interactions. Annu. Rev. Cell Dev. Biol. 39, annurev-cellbio-020223-031019. 10.1146/annurev-cellbio-020223-031019.

10. Dalghi, M.G., Montalbetti, N., Carattino, M.D., and Apodaca, G. (2020). The Urothelium: Life in a Liquid Environment. Physiol. Rev. 100, 1621–1705. 10.1152/physrev.00041.2019.

11. Jafari, N.V., and Rohn, J.L. (2022). The urothelium: a multi-faceted barrier against a harsh environment. Mucosal Immunol. 15, 1127–1142. 10.1038/s41385-022-00565-0.

12. Brigaud, I., Duteyrat, J.-L., Chlasta, J., Le Bail, S., Couderc, J.-L., and Grammont, M. (2015). Transforming Growth Factor β/activin signalling induces epithelial cell flattening during *Drosophila* oogenesis. Biol. Open 4, 345–354. 10.1242/bio.201410785.

13. Balaji, R., Weichselberger, V., and Classen, A.-K. (2019). Response of epithelial cell and tissue shape to external forces *in vivo*. Development, dev.171256. 10.1242/dev.171256.

14. Jia, D., Jevitt, A., Huang, Y.-C., Ramos, B., and Deng, W.-M. (2022). Developmental regulation of epithelial cell cuboidal-to-squamous transition in Drosophila follicle cells. Dev. Biol. 491, 113–125. 10.1016/j.ydbio.2022.09.001.

15. Pope, K.L., and Harris, T.J.C. (2008). Control of cell flattening and junctional remodeling during squamous epithelial morphogenesis in *Drosophila*. Development 135, 2227–2238. 10.1242/dev.019802.

16. Moreno-Mármol, T., Ledesma-Terrón, M., Tabanera, N., Martin-Bermejo, M.J., Cardozo, M.J., Cavodeassi, F., and Bovolenta, P. (2021). Stretching of the retinal pigment epithelium contributes to zebrafish optic cup morphogenesis. eLife 10, e63396. 10.7554/eLife.63396.

17. Grego-Bessa, J., Bloomekatz, J., Castel, P., Omelchenko, T., Baselga, J., and Anderson, K.V. (2016). The tumor suppressor PTEN and the PDK1 kinase regulate formation of the columnar neural epithelium. eLife 5, e12034. 10.7554/eLife.12034.

18. Widmann, T.J., and Dahmann, C. (2009). Dpp signaling promotes the cuboidal-to-columnar shape transition of Drosophila wing disc epithelia by regulating Rho1. J. Cell Sci. 122, 1362–1373. 10.1242/jcs.044271.

19. Harmansa, S., Erlich, A., Eloy, C., Zurlo, G., and Lecuit, T. (2023). Growth anisotropy of the extracellular matrix shapes a developing organ. Nat. Commun. 14, 1220. 10.1038/s41467-023-36739-y.

20. Kondo, T., and Hayashi, S. (2015). Mechanisms of cell height changes that mediate epithelial invagination. Dev. Growth Differ. 57, 313–323. 10.1111/dgd.12224.

21. Burnside, B. (1973). Microtubules and Microfilaments in Amphibian Neurulation. Am. Zool. 13, 989–1006. 10.1093/icb/13.4.989.

22. Tripura, C., Chandrika, N., Susmitha, V.-N., Noselli, S., and Shashidhara, L.S. (2011). Regulation and activity of JNK signaling in the wing disc peripodial membrane during adult morphogenesis in Drosophila. Int. J. Dev. Biol. 55, 583–590. 10.1387/ijdb.103275ct.

23. Halder, S., Ghosh, G., Gayen, B., and Prasad, M. (2022). TOR signalling regulates epithelial cell shape transition in *Drosophila* oogenesis (Developmental Biology) 10.1101/2022.12.29.522192.

24. Friesen, S., and Hariharan, I.K. (2023). Coordinated growth of linked epithelia is mediated by the Hippo pathway (Developmental Biology) 10.1101/2023.02.26.530099.

25. Baena-López, L.A., Pastor-Pareja, J.C., and Resino, J. (2003). Wg and Egfr signalling antagonise the development of the peripodial epithelium in *Drosophila* wing discs. Development 130, 6497–6506. 10.1242/dev.00884.

26. McClure, K.D., and Schubiger, G. (2005). Developmental analysis and squamous morphogenesis of the peripodial epithelium in Drosophila imaginal discs. Development 132, 5033–5042. 10.1242/dev.02092.

27. Nusinow, D., Greenberg, L., and Hatini, V. (2008). Reciprocal roles for *bowl* and *lines* in specifying the peripodial epithelium and the disc proper of the *Drosophila* wing primordium. Development 135, 3031–3041. 10.1242/dev.020800.

28. Tang, W., Wang, D., and Shen, J. (2016). Asymmetric distribution of Spalt in Drosophila wing squamous and columnar epithelia ensures correct cell morphogenesis. Sci. Rep. 6, 30236. 10.1038/srep30236.

29. Bonche, R., Smolen, P., Chessel, A., Boisivon, S., Pisano, S., Voigt, A., Schaub, S., Thérond, P., and Pizette, S. (2022). Regulation of the collagen IV network by the basement membrane protein perlecan is crucial for squamous epithelial cell morphogenesis and organ architecture. Matrix Biol., S0945053X22001263. 10.1016/j.matbio.2022.10.004.

30. Zatulovskiy, E., and Skotheim, J.M. (2020). On the Molecular Mechanisms Regulating Animal Cell Size Homeostasis. Trends Genet. 36, 360–372. 10.1016/j.tig.2020.01.011.

31. Balachandra, S., Sarkar, S., and Amodeo, A.A. (2022). The Nuclear-to-Cytoplasmic Ratio: Coupling DNA Content to Cell Size, Cell Cycle, and Biosynthetic Capacity. Annu. Rev. Genet. 56, 165–185. 10.1146/annurev-genet-080320-030537.

32. Neurohr, G.E., Terry, R.L., Lengefeld, J., Bonney, M., Brittingham, G.P., Moretto, F., Miettinen, T.P., Vaites, L.P., Soares, L.M., Paulo, J.A., et al. (2019). Excessive Cell Growth Causes Cytoplasm Dilution And Contributes to Senescence. Cell 176, 1083–1097.e18. 10.1016/j.cell.2019.01.018.

33. Øvrebø, J.I., and Edgar, B.A. (2018). Polyploidy in tissue homeostasis and regeneration. Development 145, dev156034. 10.1242/dev.156034.

34. Nematbakhsh, A., Levis, M., Kumar, N., Chen, W., Zartman, J.J., and Alber, M. (2020). Epithelial organ shape is generated by patterned actomyosin contractility and maintained by the extracellular matrix. PLOS Comput. Biol. 16, e1008105. 10.1371/journal.pcbi.1008105.

35. Sui, L., Pflugfelder, G.O., and Shen, J. (2012). The Dorsocross T-box transcription factors promote tissue morphogenesis in the *Drosophila* wing imaginal disc. Development 139, 2773–2782. 10.1242/dev.079384.

36. Sui, L., Alt, S., Weigert, M., Dye, N., Eaton, S., Jug, F., Myers, E.W., Jülicher, F., Salbreux, G., and Dahmann, C. (2018). Differential lateral and basal tension drive folding of Drosophila wing discs through two distinct mechanisms. Nat. Commun. 9, 4620. 10.1038/s41467-018-06497-3.

37. Diaz-de-la-Loza, M.-C., Ray, R.P., Ganguly, P.S., Alt, S., Davis, J.R., Hoppe, A., Tapon, N., Salbreux, G., and Thompson, B.J. (2018). Apical and Basal Matrix Remodeling Control Epithelial Morphogenesis. Dev. Cell 46, 23–39.e5. 10.1016/j.devcel.2018.06.006.

38. Lee, T., and Luo, L. (1999). Mosaic Analysis with a Repressible Cell Marker for Studies of Gene Function in Neuronal Morphogenesis. Neuron 22, 451–461. 10.1016/S0896-6273(00)80701-1.

39. Wilkin, M.B., Becker, M.N., Mulvey, D., Phan, I., Chao, A., Cooper, K., Chung, H.-J., Campbell, I.D., Baron, M., and MacIntyre, R. (2000). Drosophila Dumpy is a gigantic extracellular protein required to maintain tension at epidermal–cuticle attachment sites. Curr. Biol. 10, 559–567. 10.1016/S0960-9822(00)00482-6.

40. Etournay, R., Popović, M., Merkel, M., Nandi, A., Blasse, C., Aigouy, B., Brandl, H., Myers, G., Salbreux, G., Jülicher, F., et al. (2015). Interplay of cell dynamics and epithelial tension during morphogenesis of the Drosophila pupal wing. eLife 4, e07090. 10.7554/eLife.07090.

41. Ray, R.P., Matamoro-Vidal, A., Ribeiro, P.S., Tapon, N., Houle, D., Salazar-Ciudad, I., and Thompson, B.J. (2015). Patterned Anchorage to the Apical Extracellular Matrix Defines Tissue Shape in the Developing Appendages of Drosophila. Dev. Cell 34, 310–322. 10.1016/j.devcel.2015.06.019.

42. Chu, W.-C., and Hayashi, S. (2021). Mechano-chemical enforcement of tendon apical ECM into nano-filaments during Drosophila flight muscle development. Curr. Biol. 31, 1366–1378.e7. 10.1016/j.cub.2021.01.010.

43. Ayukawa, T., Akiyama, M., Hozumi, Y., Ishimoto, K., Sasaki, J., Senoo, H., Sasaki, T., and Yamazaki, M. (2022). Tissue flow regulates planar cell polarity independently of the Frizzled core pathway. Cell Rep. 40, 111388. 10.1016/j.celrep.2022.111388.

44. Plaza, S., Chanut-Delalande, H., Fernandes, I., Wassarman, P.M., and Payre, F. (2010). From A to Z: apical structures and zona pellucida-domain proteins. Trends Cell Biol. 20, 524–532. 10.1016/j.tcb.2010.06.002.

45. Jovine, L., Darie, C.C., Litscher, E.S., and Wassarman, P.M. (2005). ZONA PELLUCIDA DOMAIN PROTEINS. Annu. Rev. Biochem. 74, 83–114. 10.1146/annurev.biochem.74.082803.133039.

46. Diaz-de-la-Loza, M.-C., Loker, R., Mann, R.S., and Thompson, B.J. (2020). Control of tissue morphogenesis by the HOX gene *Ultrabithorax*. Development 147, dev184564. 10.1242/dev.184564.

47. Lowe, N., Rees, J.S., Roote, J., Ryder, E., Armean, I.M., Johnson, G., Drummond, E., Spriggs, H., Drummond, J., Magbanua, J.P., et al. (2014). Analysis of the expression patterns, subcellular localisations and interaction partners of *Drosophila* proteins using a *pigP* protein trap library. Development 141, 3994–4005. 10.1242/dev.111054.

48. Lye, C.M., Naylor, H.W., and Sanson, B. (2014). Subcellular localisations of the CPTI collection of YFP-tagged proteins in *Drosophila* embryos. Development 141, 4006–4017. 10.1242/dev.111310.

49. Nellen, D., Burke, R., Struhl, G., and Basler, K. (1996). Direct and Long-Range Action of a DPP Morphogen Gradient. Cell 85, 357–368. 10.1016/S0092-8674(00)81114-9.

50. Martín-Castellanos, C., and Edgar, B.A. (2002). A characterization of the effects of Dpp signaling on cell growth and proliferation in the *Drosophila* wing. Development 129, 1003– 1013. 10.1242/dev.129.4.1003.

51. Kolahi, K.S., White, P.F., Shreter, D.M., Classen, A.-K., Bilder, D., and Mofrad, M.R.K. (2009). Quantitative analysis of epithelial morphogenesis in Drosophila oogenesis: New insights based on morphometric analysis and mechanical modeling. Dev. Biol. 331, 129–139. 10.1016/j.ydbio.2009.04.028.

52. Leiby, K.L., Yuan, Y., Ng, R., Raredon, M.S.B., Adams, T.S., Baevova, P., Greaney, A.M., Hirschi, K.K., Campbell, S.G., Kaminski, N., et al. (2023). Rational engineering of lung alveolar epithelium. Npj Regen. Med. 8, 22. 10.1038/s41536-023-00295-2.

53. Drees, L., Schneider, S., Riedel, D., Schuh, R., and Behr, M. (2023). The proteolysis of ZP proteins is essential to control cell membrane structure and integrity of developing tracheal tubes in Drosophila. eLife 12, e91079. 10.7554/eLife.91079.

54. Haltom, A.R., and Jafar-Nejad, H. (2015). The multiple roles of epidermal growth factor repeat *O* -glycans in animal development. Glycobiology 25, 1027–1042. 10.1093/glycob/cwv052.

55. Strigini, M., and Cohen, S.M. (2000). Wingless gradient formation in the Drosophila wing. Curr. Biol. 10, 293–300. 10.1016/S0960-9822(00)00378-X.

56. Kalukula, Y., Stephens, A.D., Lammerding, J., and Gabriele, S. (2022). Mechanics and functional consequences of nuclear deformations. Nat. Rev. Mol. Cell Biol. 23, 583–602. 10.1038/s41580-022-00480-z.

57. Nader, G.P.D.F., Williart, A., and Piel, M. (2021). Nuclear deformations, from signaling to perturbation and damage. Curr. Opin. Cell Biol. 72, 137–145. 10.1016/j.ceb.2021.07.008.

58. Gong, J., Nirala, N.K., Chen, J., Wang, F., Gu, P., Wen, Q., Ip, Y.T., and Xiang, Y. (2023). TrpA1 is a shear stress mechanosensing channel regulating intestinal stem cell proliferation in *Drosophila*. Sci. Adv. 9, eadc9660. 10.1126/sciadv.adc9660.

59. Fletcher, G.C., Diaz-de-la-Loza, M.-C., Borreguero-Muñoz, N., Holder, M., Aguilar-Aragon, M., and Thompson, B.J. (2018). Mechanical strain regulates the Hippo pathway in *Drosophila*. Development 145, dev159467. 10.1242/dev.159467.

60. Tanimoto, H., Itoh, S., Ten Dijke, P., and Tabata, T. (2000). Hedgehog Creates a Gradient of DPP Activity in Drosophila Wing Imaginal Discs. Mol. Cell 5, 59–71. 10.1016/S1097-2765(00)80403-7.

61. Morin, X., Daneman, R., Zavortink, M., and Chia, W. (2001). A protein trap strategy to detect GFP-tagged proteins expressed from their endogenous loci in Drosophila. Proc. Natl. Acad. Sci. 98, 15050–15055. 10.1073/pnas.261408198.

62. Tsuneizumi, K., Nakayama, T., Kamoshida, Y., Kornberg, T.B., Christian, J.L., and Tabata, T. (1997). Daughters against dpp modulates dpp organizing activity in Drosophila wing development. Nature 389, 627–631. 10.1038/39362.

63. Harmansa, S., Hamaratoglu, F., Affolter, M., and Caussinus, E. (2015). Dpp spreading is required for medial but not for lateral wing disc growth. Nature 527, 317–322. 10.1038/nature15712.

64. Dye, N.A., Popović, M., Spannl, S., Etournay, R., Kainmüller, D., Ghosh, S., Myers, E.W., Jülicher, F., and Eaton, S. (2017). Cell dynamics underlying oriented growth of the *Drosophila* wing imaginal disc. Development, dev.155069. 10.1242/dev.155069.

65. Van De Bor, V., Loreau, V., Malbouyres, M., Cerezo, D., Placenti, A., Ruggiero, F., and Noselli, S. (2021). A dynamic and mosaic basement membrane controls cell intercalation in *Drosophila* ovaries. Development 148, dev195511. 10.1242/dev.195511.

66. Herbert, S., Valon, L., Mancini, L., Dray, N., Caldarelli, P., Gros, J., Esposito, E., Shorte, S.L., Bally-Cuif, L., Aulner, N., et al. (2021). LocalZProjector and DeProj: a toolbox for local 2D projection and accurate morphometrics of large 3D microscopy images. BMC Biol. 19, 136. 10.1186/s12915-021-01037-w.

67. Aigouy, B., Umetsu, D., and Eaton, S. (2016). Segmentation and Quantitative Analysis of Epithelial Tissues. In Drosophila Methods in Molecular Biology., C. Dahmann, ed. (Springer New York), pp. 227–239. 10.1007/978-1-4939-6371-3_13.

68. Waskom, M. (2021). seaborn: statistical data visualization. J. Open Source Softw. 6, 3021. 10.21105/joss.03021.

69. Mary, H., and Brouhard, G.J. (2019). Kappa ( *κ* ): Analysis of Curvature in Biological Image Data using B-splines (Bioinformatics) 10.1101/852772.

70. Berg, S., Kutra, D., Kroeger, T., Straehle, C.N., Kausler, B.X., Haubold, C., Schiegg, M., Ales, J., Beier, T., Rudy, M., et al. (2019). ilastik: interactive machine learning for (bio)image analysis. Nat. Methods 16, 1226–1232. 10.1038/s41592-019-0582-9.

71. Tasnadi, E.A., Toth, T., Kovacs, M., Diosdi, A., Pampaloni, F., Molnar, J., Piccinini, F., and Horvath, P. (2020). 3D-Cell-Annotator: an open-source active surface tool for single-cell segmentation in 3D microscopy images. Bioinformatics 36, 2948–2949. 10.1093/bioinformatics/btaa029.

72. Dear, M.L., Dani, N., Parkinson, W., Zhou, S., and Broadie, K. (2015). Two matrix metalloproteinase classes reciprocally regulate synaptogenesis. Development, dev.124461. 10.1242/dev.124461.

73. Schindelin, J., Arganda-Carreras, I., Frise, E., Kaynig, V., Longair, M., Pietzsch, T., Preibisch, S., Rueden, C., Saalfeld, S., Schmid, B., et al. (2012). Fiji: an open-source platform for biological-image analysis. Nat. Methods 9, 676–682. 10.1038/nmeth.2019.

