## Supplemental_Figures for "Mechanical regulation of cuboidal-to-squamous epithelial transition in the *Drosophila* developing wing"

### Supplementary Figures – Harmansa and Lecuit

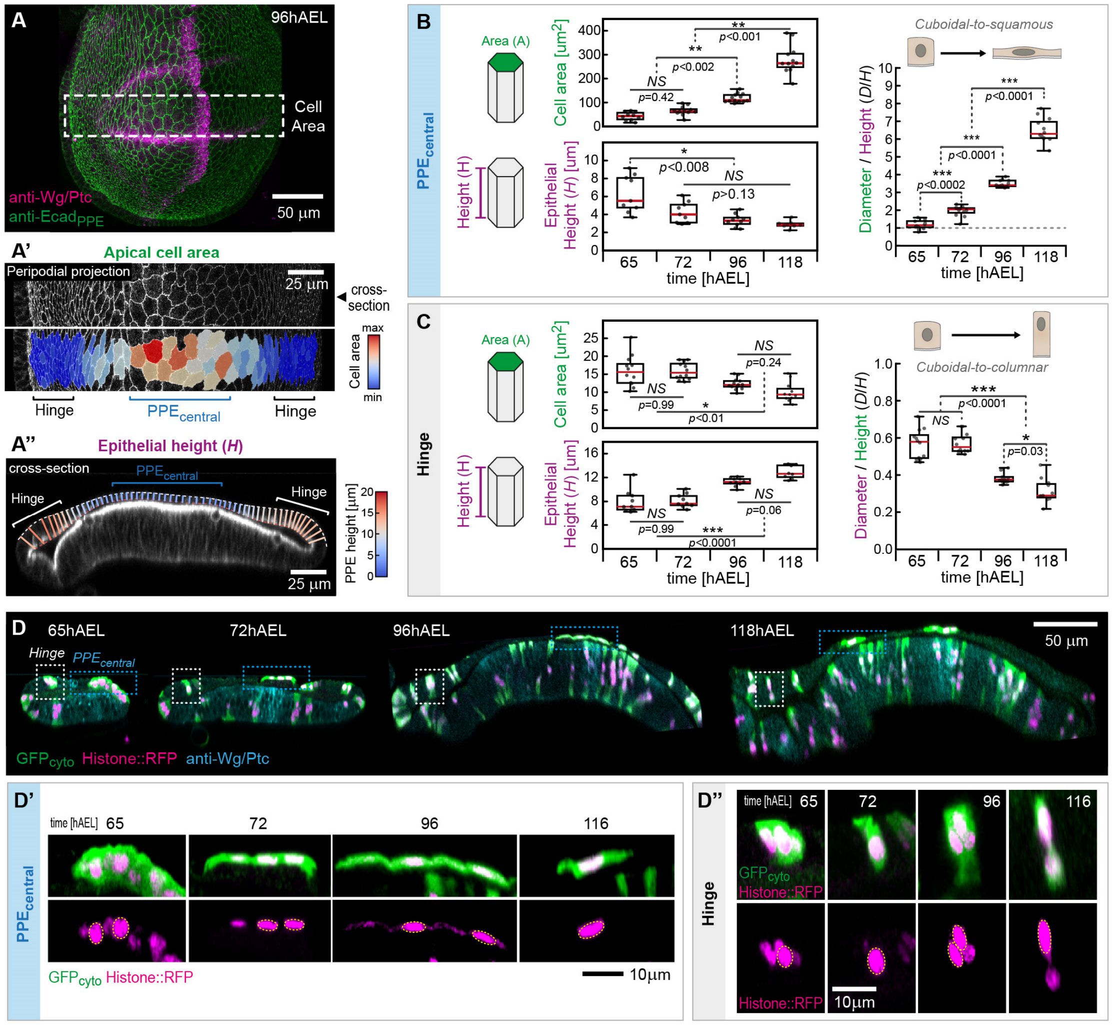


**Supplementary Figure 1 - Dynamics of cell shape transitions during peripodial morphogenesis**
(**A**) Quantification of peripodial cell area and tissue height: *top*: PPE surface projection of the $Ecad_{PPE}$ signal overlaid with the maximum projection of the Wg/Ptc signal (PPE+DP). Peripodial cell area was quantified in the region dorsal to the D/V-boundary of the DP (marked by Wg) as indicated by the dashed white box. (A') Magnification of the $Ecad_{PPE}$ in the region of interest with color-coded segmented apical cell area (maximum cell area in red, minimum cell area in blue). The domains of the $PPE_{central}$ (blue) and the peripheral hinge (black) are marked by brackets. The position of the cross-section used for height quantification is indicated. (A'') Cross-section view of disc marked for F-Actin to visualize cell outlines. Local PPE height is shown by color-code and $PPE_{central}$ (blue) and hinge region (black) are marked by brackets. (**B**+**C**) Quantification of $PPE_{central}$ cell area (*A*, green, *top left*), epithelial height (*H*, purple, *bottom left*) and cellular aspect ratio (D/H, *right*) in the $PPE_{central}$ (B, blue) and the peripodial hinge (C, gray). Sample-numbers for cell area [cells/discs]: $n_{65}$=161/12, $n_{72}$=759/12, $n_{96}$=3246/13, $n_{118}$=1812/12, epithelial height [discs]: $n_{65}$=9, $n_{72}$=9, $n_{96}$=10, $n_{118}$=8 and cellular aspect ratio [cells/discs]: $n_{65}$=180/12, $n_{72}$=758/12, $n_{96}$=3246/13, $n_{118}$=1812/12. (**D**) Optical cross-sections of wing discs at indicated time points expressing cytosolic GFP ($GFP_{cyto}$, green) and Histone::RFP (magenta) in a clonal manner. Magnifications of example clones of central PPE ($PPE_{central}$) and hinge cells are shown in (D') and (D''), respectively. *Statistics*: Statistical significance was assessed by a one-way ANOVA and Tukey's post hoc test ( $*p \leq 0.05$, $**p \leq 0.005$, $***p \leq 0.0005$). In box plots, the median is indicated by a central thick line, while a box outlines the interquartile range (containing 50% of the data points). Whiskers indicate the minimum and maximum data range.

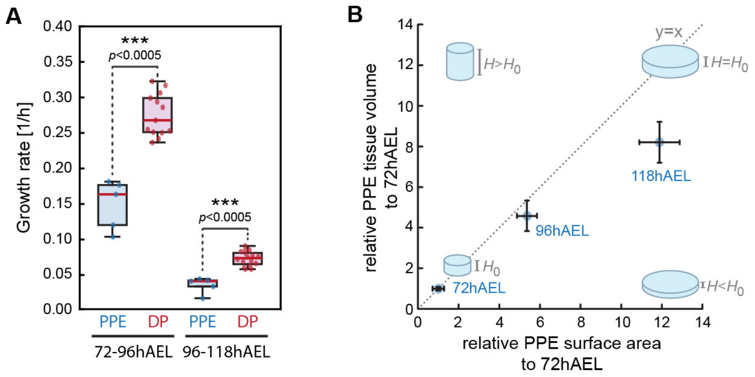


*Data from Harmansa et.al. (2023) Nature Communications*
Supplementary Figure 1k

**Supplementary Figure 2 – PPE and DP growth dynamics**
(**A**) Growth rate in the PPE and the DP between 72-96hAEL and 96-118hAEL. The growth rate was calculated using the volume date for the PPE and DP from Supplementary Figure 1F in Harmansa et.al.[1]. In boxplots, the median is indicated by a central thick line, while a box outlines the interquartile range (containing 50% of the data points). Whiskers indicate the minimum and maximum data range. Statistical significance was assessed by a two-sided Student's *t*-test (unequal variance, *P ≤ 0.05, **P ≤ 0.005, ***P ≤ 0.0005). (**B**) Plot of the relative changes in the surface area covered by the PPE versus its tissue volume (quantifying the tissue covering the DP Wingless ring, data from Supplementary Figure 1 in Harmansa *et.al.*[1]). The original tissue height ($H_0$) remains unchanged if area and volume increase IS proportional (dashed line). In contrast, if the area change exceeds the volumetric growth of the PPE the tissue will be dilated/stretched leading to a decrease in tissue height ($H<H_0$). Error bars indicate the standard error. Sample numbers [discs] for area: $n_{72}$= 24, $n_{96}$ = 13, $n_{118}$ = 16 and for volume: $n_{72}$= 5, $n_{96}$ = 5, $n_{118}$ = 5.

1. Harmansa, S., Erlich, A., Eloy, C., Zurlo, G., and Lecuit, T. (2023). Growth anisotropy of the extracellular matrix shapes a developing organ. Nat. Commun. *14*, 1220. https://doi.org/10.1038/s41467-023-36739-y.

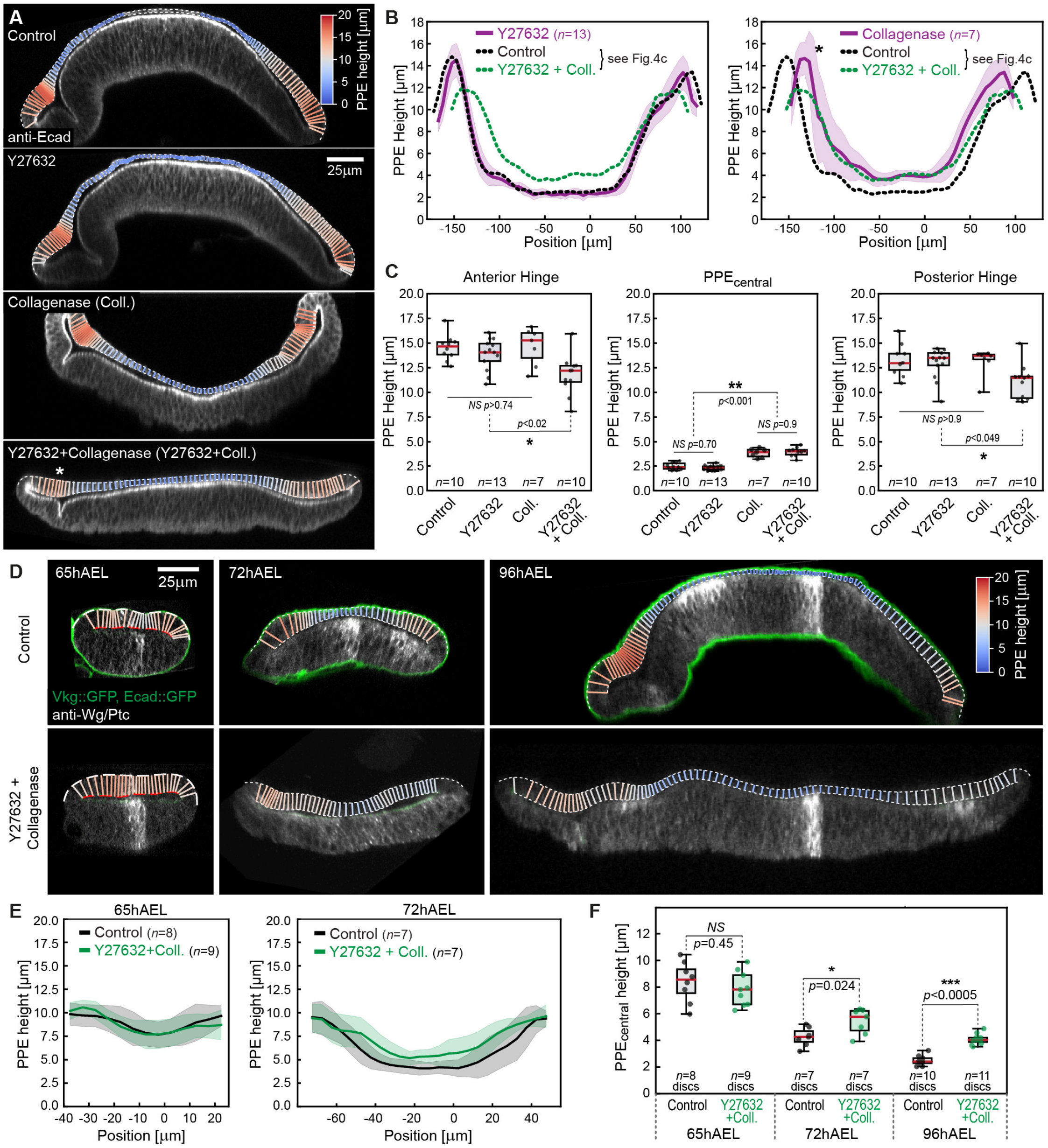


**Supplementary Figure 3 - Acute BM digestion and concomitant MyoII inhibition**

(**A**) Representative cross-sections of 96hAEL wing discs treated as indicated. PPE height is color-coded. (**B**) *left*: PPE height upon treatment with Y27632 only (magenta) compared to control (black) or Y27632+Coll. discs (green). *right*: PPE height in discs only treated with Collagenase (magenta) compared to control (black) and Y27632+Coll. discs (green). (**C**) PPE height in the anterior hinge (*left*), the central PPE (*middle*) and the posterior hinge (*right*) for indicated conditions. Sample numbers indicate the number of discs. (**D**) Section views of Vkg::GFP (green) wing discs at indicated developmental time points (65, 72 and 96hAEL) not-treated (control, *top*) and treated with Y27632 and Collagenase (*bottom*). Peripodial height is color-coded. (**E**) Quantification of PPE height in non-treated and Y27632+Collagen (Coll.) discs at 65hAEL (*left*) and at 72hAEL (*right*). *n*-numbers indicate the number of discs. (**F**) Quantification of epithelial height of the central, squamous PPE cells at indicated time points and conditions. *Statistics*: Error bands in (B+E) indicate standard deviation. Red line in box plots in (C+F) indicates the median, boxes mark the interquartile range (50% of data points) and whiskers the data range. Statistical significance was assessed by a one-way ANOVA and Tukey's post hoc test in (C) and by a two-sided Student's *t*-test in (F) (unequal variance, *P ≤ 0.05, **P ≤ 0.005, ***P ≤ 0.0005).

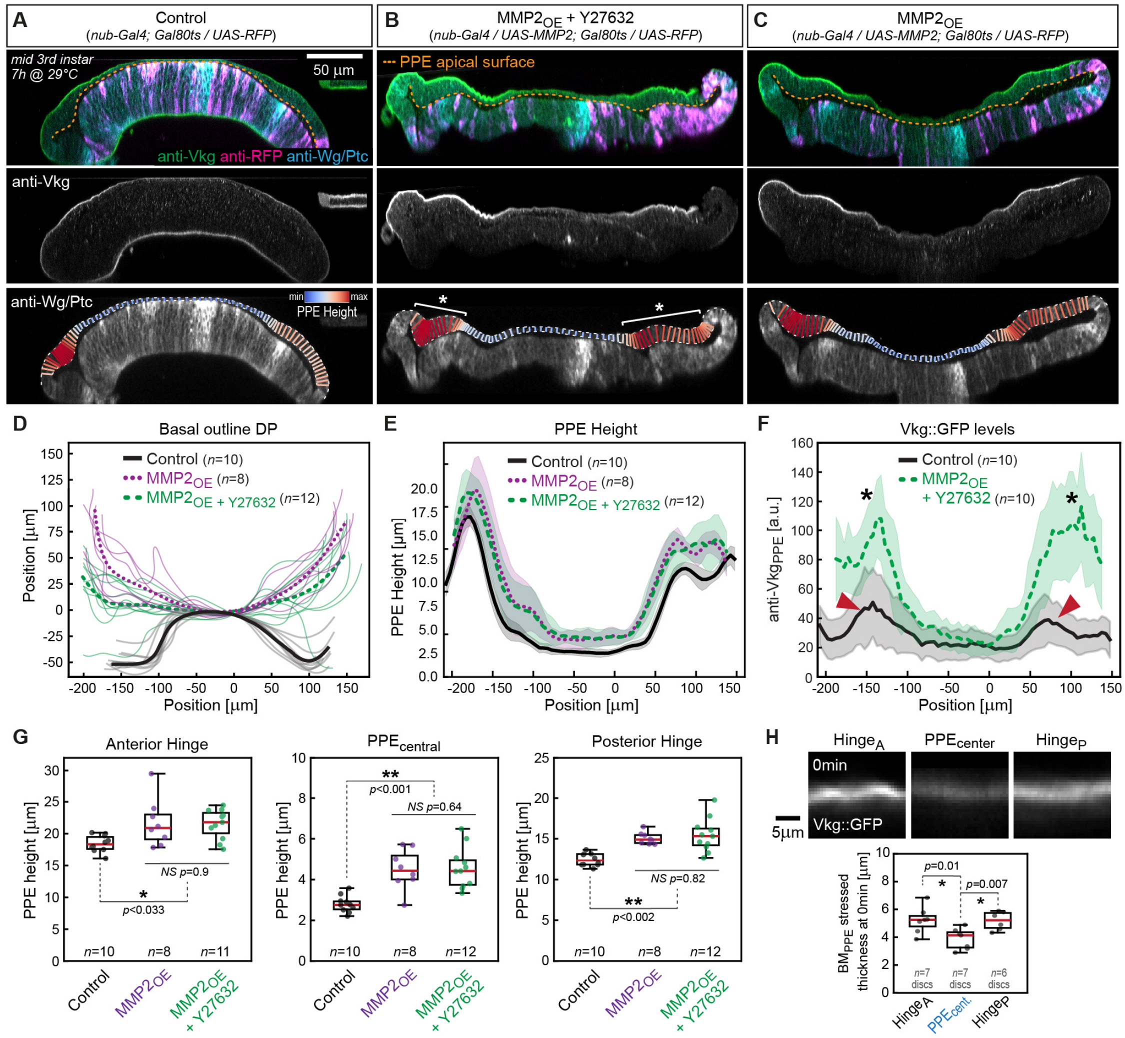


**Supplementary Figure 4 - Genetic digestion of the disc proper BM$_{DP}$**

(**A-C**) Section-views of representative mid-3[rd] instar wing discs expressing either RFP (control, (**A**)) or RFP and MMP2 (MMP2$_{OE}$, (**B**) and (**C**)) in the DP (under control of *nubbin-Gal4* (nub-Gal4)). MMP2 overexpressed was induced 7h before dissection by a shift to 29°C using the Gal80$^{ts}$ system (see methods). In addition, the disc shown in (B) was treated with the Rock inhibitor Y27632 (Myosin II inhibition) after dissection, leading to disc flattening (referred to as MMP$_{OE}$ + Y27632). Discs were stained for Wg/Ptc (disc outline) and Vkg (*middle*). PPE height is indicated by colour code (*bottom*). Ectopic MMP2 expression in the DP leads to BM degradation and a loss of disc bending. However, the hinge$_A$ and hinge$_P$ do not relax in the presence of the BMPPE (asterisks). (**D**) Average outlines of the basal DP surface of control discs (black) and discs overexpressing MMP2 (magenta) and overexpressing MMP + Y27632 treatment (green). Fine lines correspond to individual discs while thick lines represent the average outlines per condition. (**E**) Average PPE height in indicated conditions. (**F**) Peripodial Vkg intensity quantified along the A/P axis in indicated conditions. Vkg intensity is higher in the hinge than in the centre in controls (bend disc, see red arrowheads). Loss of disc bending leads to a strong increase in Vkg levels in the hinge (asterisks). (**G**) Quantification of PPE height in the anterior hinge (*left*), the central squamous cells (*middle*) and the posterior hinge (*right*) for the different conditions. (**H**) *top*: Cross sections of Vkg::GFP in the stressed configuration (0min, related to Figure 5D). *bottom*: Quantification of stressed BM$_{PPE}$ thickness. *Statistics*: *n*-numbers indicate the number of discs. Error bands in (E+F) indicate standard deviation. Red line in box plots in (G+H) indicates the median, boxes mark the interquartile range (50% of data points) and whiskers the data range. Statistical significance was assessed by a one-way ANOVA and Tukey's post hoc test ( $*p \leq 0.05$, $**p \leq 0.005$, $***p \leq 0.0005$).

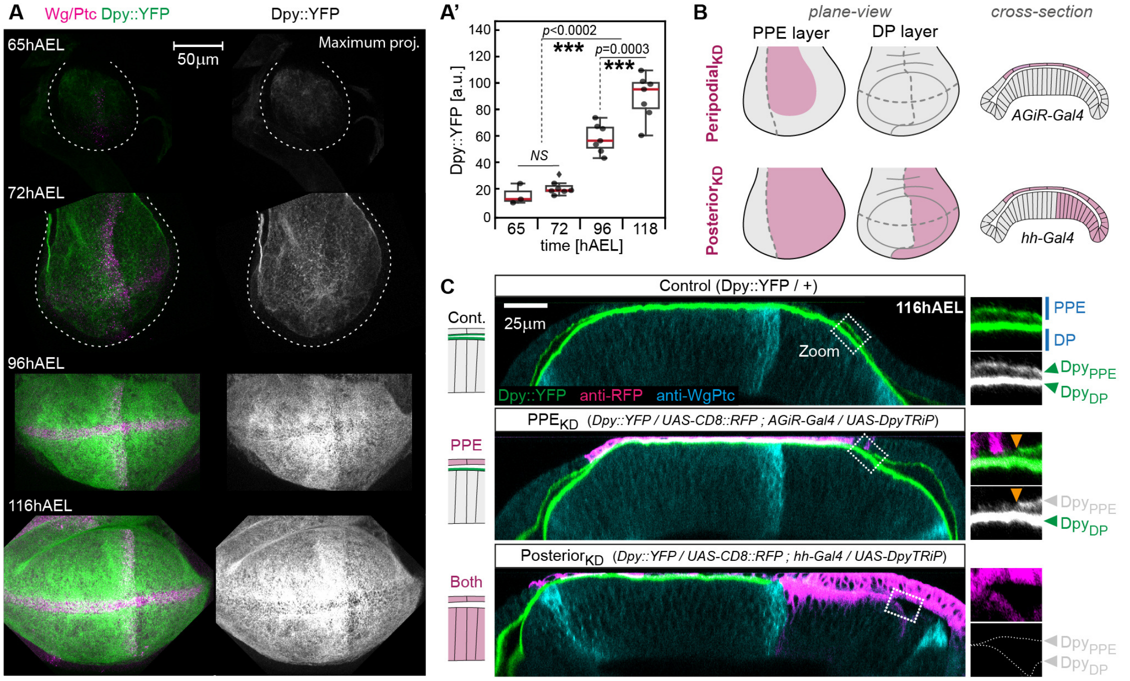


**Supplementary Figure 5 - Dpy expression during development and knock-down conditions**
(**A**) Maximum projection of representative Dpy::YFP wing discs at indicated time points, stained for YFP (green) and Wg/Ptc (magenta). In young discs, the outline of the tissue is indicated by a dashed line. Average fluorescent Dpy:YFP levels in the wing pouch area (marked by the Wg staining) are quantified in (A'). Sample numbers [discs]: $n_{65}$=3, $n_{72}$=7, $n_{96}$=7, $n_{116}$=7. (**B**) Scheme illustrating the spatial expression profiles of the used Gal4-driver lines (magenta) in the PPE- (*left*), the DP-layer (*middle*) and in section view (*right*). *AGiR-Gal4* is expressed in the central squamous cells of the PPE (*top*) and *hh-Gal4* in the posterior compartment of both, the DP- and the PPE-layer (*bottom*). (**C**) Section-view of representative wing discs expressing Dpy::YFP (green) either alone (control, *top*) or together with a Dpy TRiP-line in the PPE (*AGiR-Gal4*, PPE$_{KD}$) or in the posterior compartment (*hh-Gal4*, Posterior$_{KD}$). Regions enlarged to the right visualize the loss of either the peripodial fraction (Dpy$_{PPE}$), the disc proper fraction (Dpy$_{DP}$) or both fractions, indicated in gray. *Statistics*: *n*-numbers indicate the number of discs. Statistics: Red line in box plots in (A') indicates the median, boxes mark the interquartile range (50% of data points) and whiskers the data range. Statistical significance was assessed by a one-way ANOVA and Tukey's post hoc test ( *$p \leq 0.05$, **$p \leq 0.005$, ***$p \leq 0.0005$).

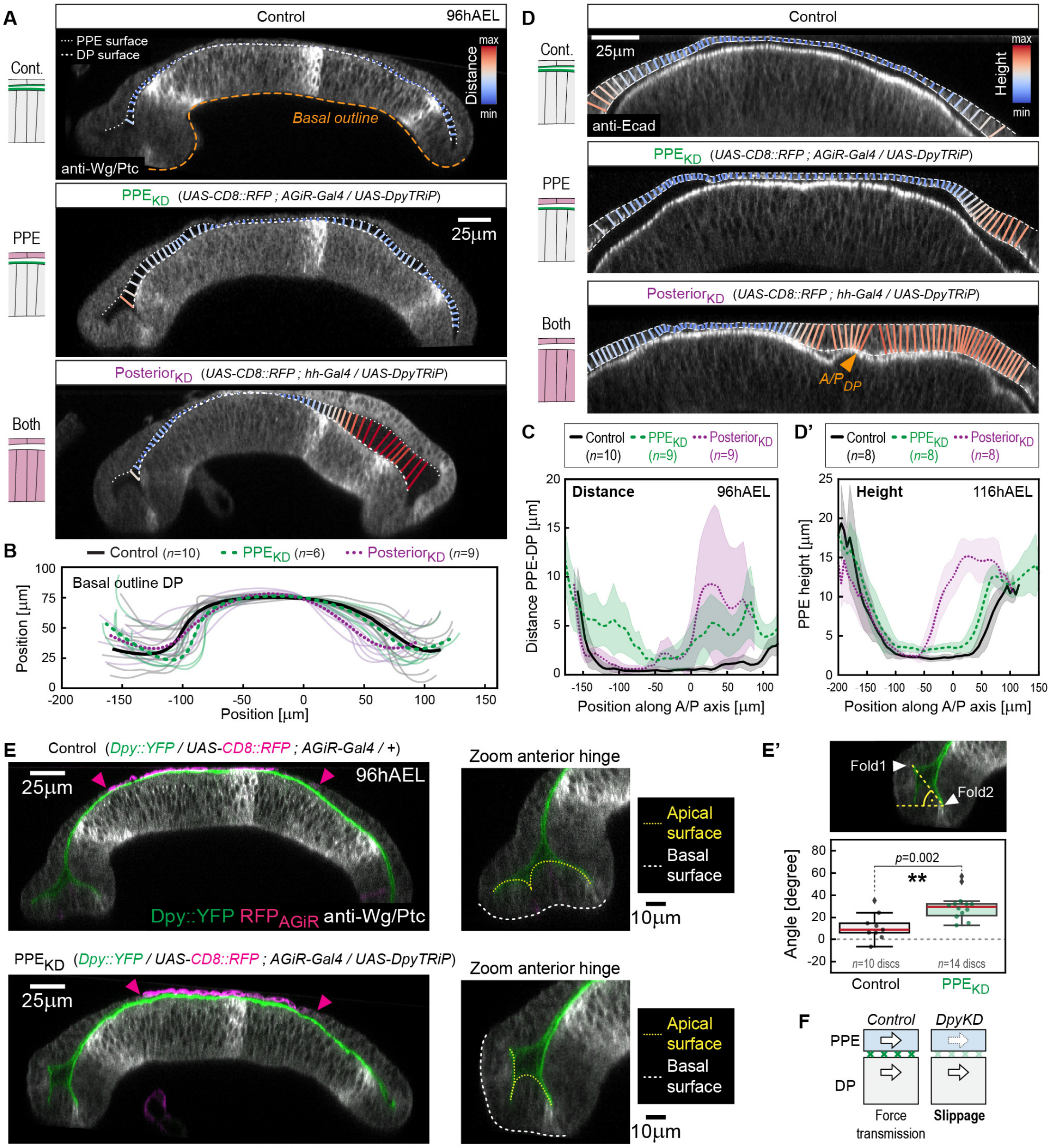


**Supplementary Figure 6 - Dpy knock-down in different regions of the growing wing disc**

(**A**) Cross section views of representative win discs of indicated genotypes. The distance between the apical surface of the DP and the PPE (marked by dashed white lines) is quantified by a color code. (**B**) Quantification of the average basal outline of the DP epithelium (as indicated in C by dashed orange line). to assess disc bending. Individual discs are shown in faint colors while the average outlines are shown by thick lines. (**C**) Quantification of the apical distance between DP and PPE in the different genotypes as indicated by color-code in (A). (**D**) Cross-sections of representative wing discs of the indicated genotypes with PPE height visualized by color-code. Continuous PPE height is quantified in (D'). (**E**) Cross sections of representative control (*top*) and PPE$_{KD}$ discs (*bottom*) expressing RFP in the squamous PPE$_{central}$ cells (magenta). The width of the AGiR expression domain is indicated by arrowheads. The anterior hinge region is magnified to the right and apical and basal surfaces are marked by yellow and white dashed lines, respectively. In PPE$_{KD}$ discs the anterior hinge is folded upwards as indicated by a positive angle between the two major Hinge$_A$ folds and the horizontal axis (E'). (**F**) Loss of Dpy (green crosses) impairs force transmission leading to slippage between the two epithelial layers and reduced shear forces acting on the PPE layer. *Statistics*: Error bands in (C+D') indicate standard deviation. Red line in box plots in (E') indicates the median, boxes mark the interquartile range (50% of data points) and whiskers the data range. Statistical significance was assessed by a two-sided Student's *t*-test (unequal variance, $^*P \leq 0.05$, $^{**}P \leq 0.005$, $^{***}P \leq 0.0005$).

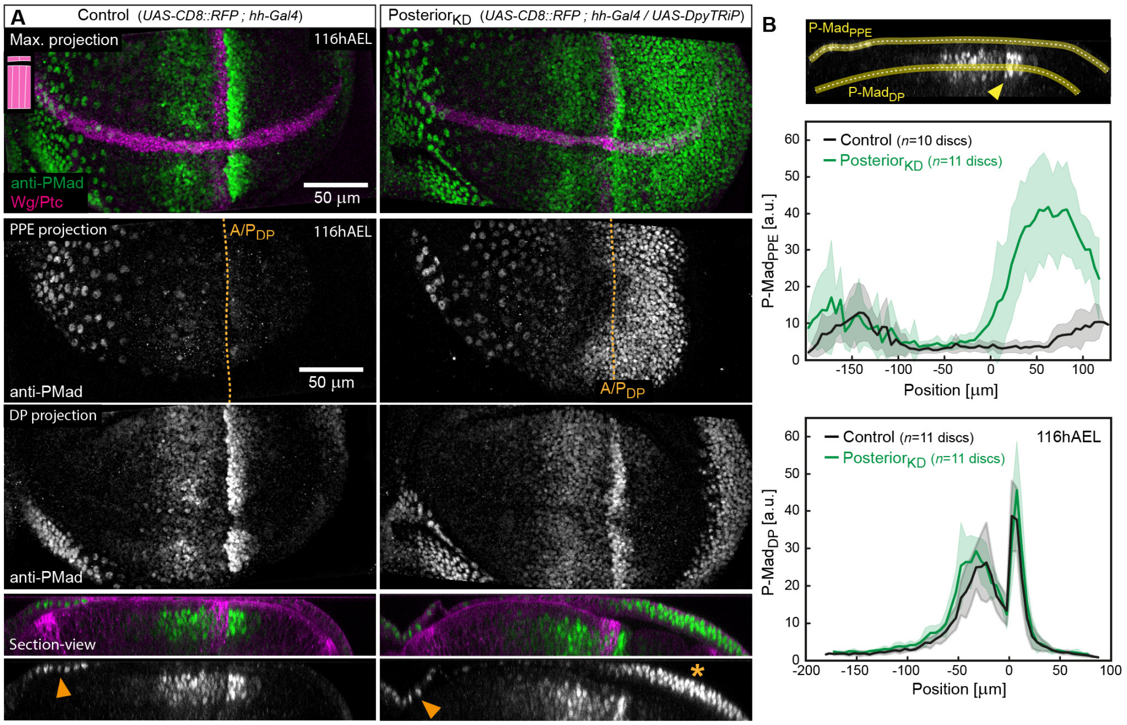


**Supplementary Figure 7 – Dpy knockdown induces peripodial Dpp signaling**

(**A**) Representative control wing disc (*left*) and posterior$_{KD}$ disc (*right*) in maximum projection (*top*), PPE projection (*middle-top*), DP projection (*middle-bottom*), and cross-section view (*bottom*) stained for the active form of the Dpp signal transducer P-Mad (green). In section views the upregulation of P-Mad due to Dpp expression at the peripodial A/P-compartment boundary is marked by an arrowhead while the ectopic posterior increase in P-Mad in Posterior$_{KD}$ discs is marked by an asterisk. (**B**) Quantification of P-Mad levels in the PPE (*top*) and DP (*bottom*) of control (black) and Posterior$_{KD}$ discs (green) at 116hAEL. P-Mad levels were quantified from cross-sections along a line as indicated in the top section view (yellow line). The A/P$_{DP}$-boundary is marked by an arrowhead and corresponds to $x$=0 in the plots. *Statistics*: Error bands in (B) indicate the standard deviation.

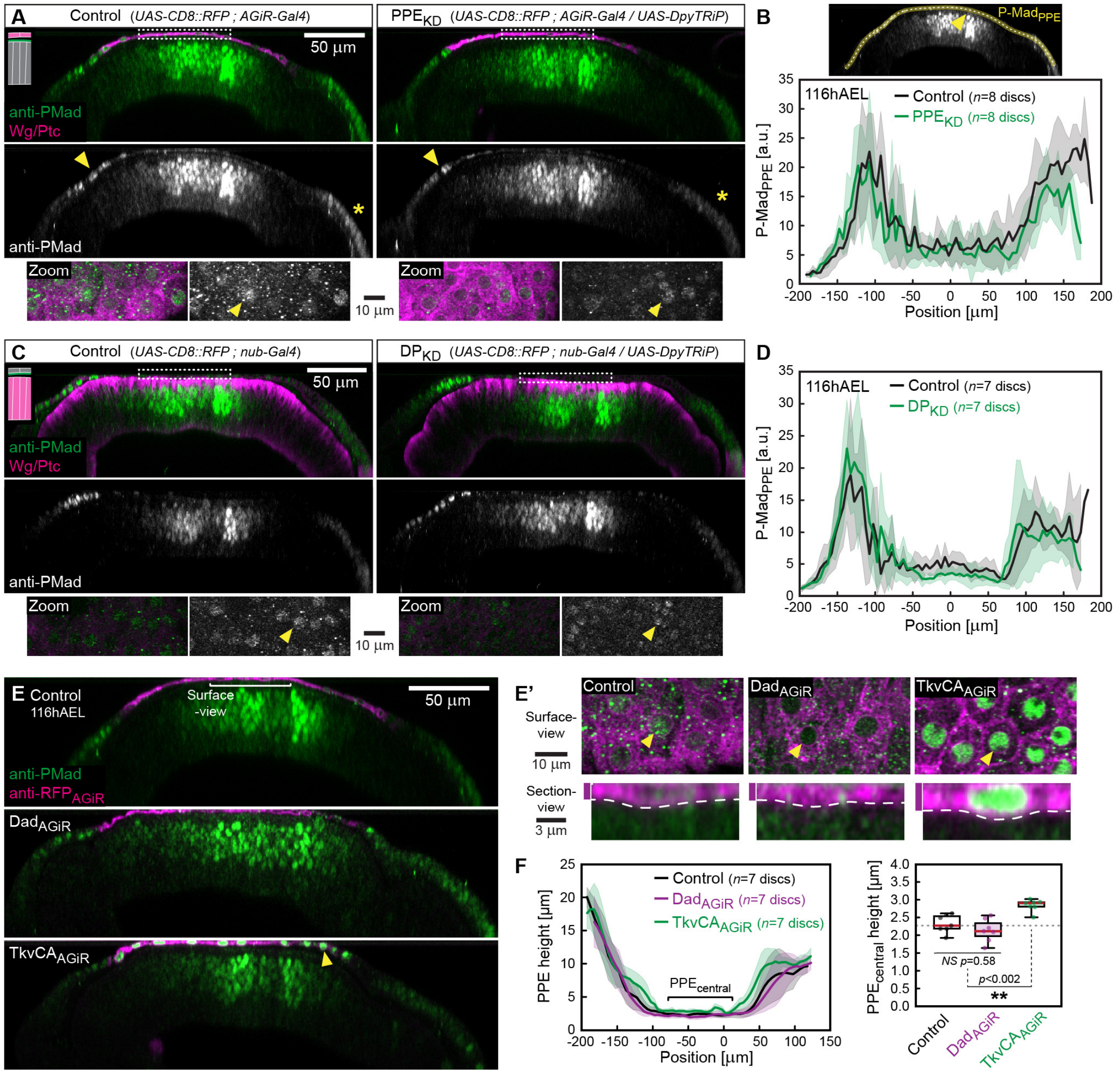


**Supplementary Figure 8 - Dpp signaling is not sufficient to increase peripodial height**

(**A**) Section-views of representative wing discs expressing CD8::mRFP alone (*left*) or together with a Dpy TRiP line (PPE$_{KD}$, *right*) in the squamous cells (*AGiR-Gal4*), stained for P-Mad (green) and mRFP (magenta). P-Mad expression induced by the A/P$_{PPE}$-boundary and posterior Dpp source are marked by an arrowhead and an asterisk, respectively. *bottom*: Zooms show PPE surface projection of the domains indicated by dashed boxes. In both conditions, low P-Mad levels are observed in squamous cells (see arrowheads). (**B**) Quantification of average P-Mad levels in the PPE layer (P-Mad$_{PPE}$, extracted along a yellow line from cross-section images). (**C-D**) As in (A-B) for wing discs in which Dpy was knocked down only in the DP layer using *nub-Gal4*, referred to as DP$_{KD}$. (**E**) Section view of wing discs expressing CD8::mRFP alone (control, *top*) or together with Dad (a suppressor of Dpp signaling, Dad$_{AGiR}$, *middle*) or with a constitutive-active form of the Dpp receptor Thickveins (TkvCA$_{AGiR}$) under the control of *AGiR-Gal4* (in squamous PPE cells, *bottom*), stained for P-Mad (green) and mRFP (magenta). Surface- and section-views of the different genotypes are shown in (E'). In (E') arrowheads mark example nuclei that show low P-Mad in control (*left*), a loss of P-Mad in Dad (*middle*), and increased P-Mad in TkvCA (*right*). A dashed line marks the apical surface of the PPE layer in section views. (**F**) Quantification of PPE height along the A/P axis (*left*) and in the squamous PPE$_{central}$ cells (*right*). *Statistics*: Error bands in (B+D+F) indicate standard deviation. Red line in box plots in (F) indicates the median, boxes mark the interquartile range (50% of data points) and whiskers the data range. Statistical significance was assessed by a one-way Anova and Tukey's post hoc test ( *$p \le 0.05$, **$p \le 0.005$, ***$p \le 0.0005$).

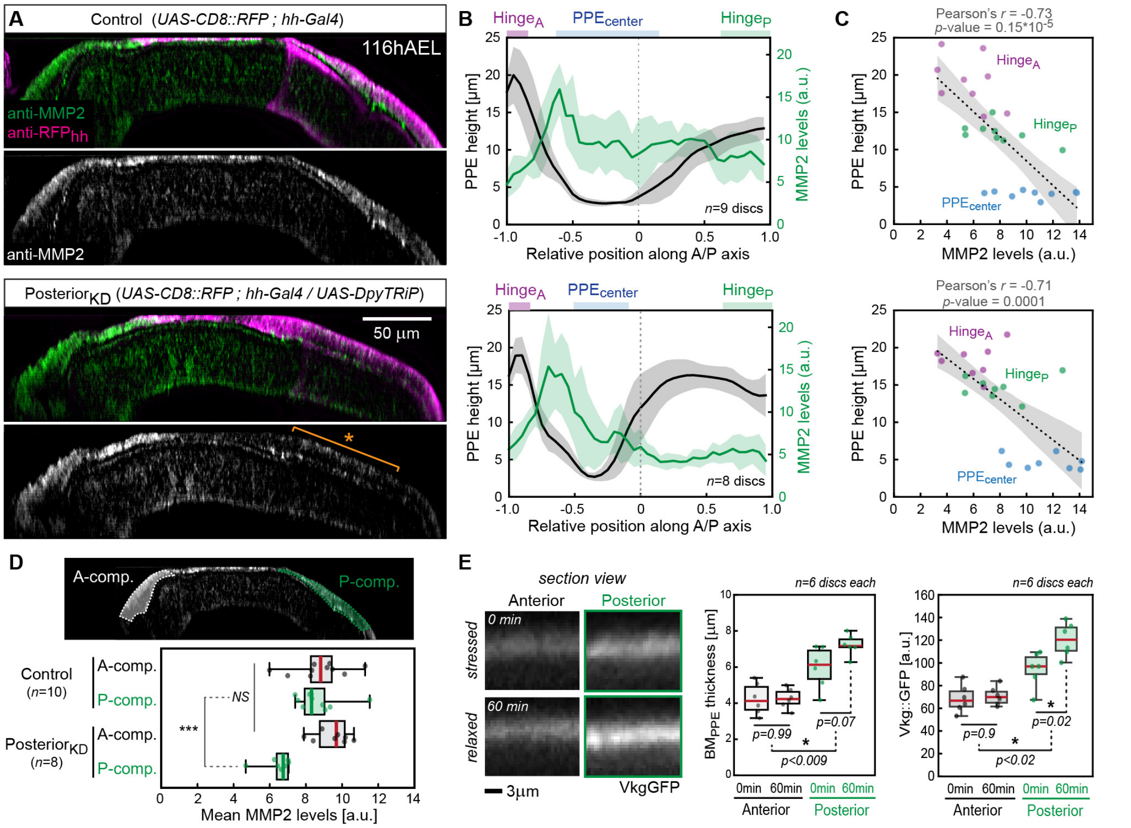


**Supplementary Figure 9 - Dpy is required for peripodial MMP2 expression**

(**A**) Cross-sections of representative wing discs expressing only CD8::mCherry in the posterior compartment (*top*, control) or together with a Dpy-TRiP line (*bottom*, $Posterior_{KD}$) stained from MMP2 (green). The posterior domain in which Dpy was knocked down shows reduced MMP2 levels, indicated by an asterisk in the $posterior_{KD}$ disc. (**B**) Quantification of MMP2 levels (green) and PPE height (grey) along the A/P axis in control (*top*) and $Posterior_{KD}$ (*bottom*). The $Hinge_A$ (magenta), $PPE_{center}$ (blue) and $Hinge_P$ (green) domains used for the quantifications shown in (C) are indicated on the top of the plots. (**C**) Plots of average MMP2 levels versus average PPE height in control (*top*) and $Posterior_{KD}$ discs (*bottom*). Spot mark average values for individual wing discs quantified in the regions indicated in (B). A linear regression is shown (dashed line) and Pearson's *r* and *p*-value are indicated. (**D**) Anti-MMP2 mean fluorescence intensities, quantified from cross-section views in the regions indicated on the scheme to the *left*. (**E**) *left*: Cross-section views of the $BM_{PPE}$ (Vkg::GFP) in $Posterior_{KD}$ discs in stressed (*top*) and relaxed configuration (*bottom*, related to Figure 7F). Quantification of $BM_{PPE}$ thickness (*middle*) and Vkg::GFP intensity (*left*) in the indicated conditions. *Statistics*: Error bands in (B) indicate standard deviation. In regression plots (C) a linear regression is makred by a dashed line and the 95% confidence interval by a grey shade. Pearson's r and p-value were calculated using the Python scipy.stats package (linregress function).In boxplots (D+E) a central thick line indicates the median, boxes mark the interquartile range (50% of data points) and whiskers the data range and statistical significance was assessed by a one-way Anova and Tukey's post hoc test ( $*p \leq 0.05$, $**p \leq 0.005$, $***p \leq 0.0005$).

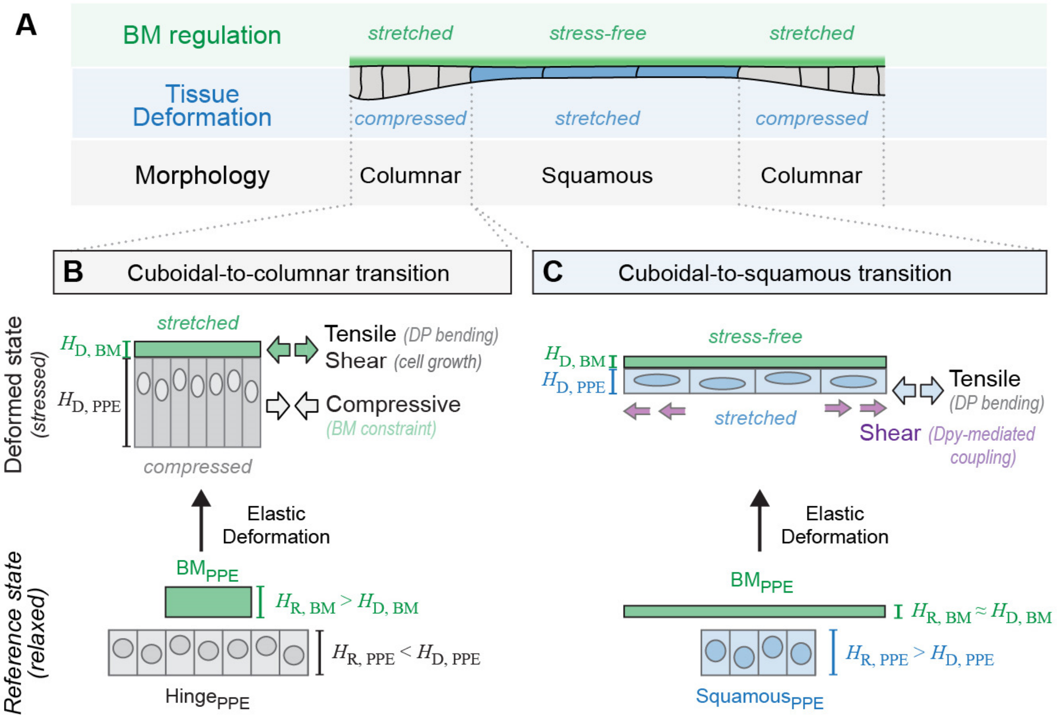


**Supplementary Figure 10 – Elastic stresses lead to deformations of the cells and BM layers driving distinct epithelial shape transitions**

(**A**) Scheme illustrating the interplay of elastic stresses accumulated in the BM$_{PPE}$ (green) and in the peripodial tissue layer (blue) leading to deformations that shape epithelial morphology (grey). (**B**) Cuboidal-to-columnar transitions in the peripodial hinge of the wing disc are the result of compressive stress in the tissue layer (deformed state, *top*). Compressive stress arises due to the hinge BM$_{PPE}$ opposing planar hinge expansion and growth (see Figure 4 and Figure S4). As a result, the unstressed, relaxed height of the hinge tissue (H$_{R,\ PPE}$, *bottom*) is lower than in the deformed state (H$_{D,\ PPE}$), where compressive stress leads to epithelial thickening. Notably, the hinge BMPPE is dual stressed: (1) by the expanding hinge tissue and (2) due to DP bending, resulting in elastic tensile stress in the hinge BM$_{PPE}$ (hence, H$_{R,\ BM}$ > H$_{R,\ BM}$, see Figure 5D and Figure S4H). (**C**) In contrast, cuboidal-to-squamous transitions result from cell stretching (deformed state, *top*) due to tensile stress and shear stress originating from DP bending (Figure 4D-E) and Dpy-mediated coupling (Figure 6D), respectively. Notably, tissue stretching depends on the central BM$_{PPE}$ to be stress-free (Figure 5C) allowing the tensile and shear stress to act on and flatten the cell layer. The stress-free state of the BM$_{PPE}$ depends on planar BM$_{PPE}$ growth (i.e. H$_{R,\ BM}$ ≈ H$_{R,\ BM}$) as any deviation of planar BM$_{PPE}$ growth leads to tensile stress in the BM$_{PPE}$, shielding the tissue layer form stretching (Figure 7C-F).
